# Lipid Droplets Promote Phase Separation of Ago2 to Accelerate Dicer1 Loss and Decelerate miRNA Activity in Lipid Exposed Hepatic Cells

**DOI:** 10.1101/2020.11.30.405449

**Authors:** Diptankar Bandyopadhyay, Sudarshana Basu, Ishita Mukherjee, Ritobrita Chakraborty, Kamalika Mukherjee, Krishnananda Chattopadhyay, Saikat Chakrabarti, Partha Chakrabarti, Suvendra N. Bhattacharyya

**Affiliations:** RNA Biology Research Laboratory, Molecular Genetics Division, CSIR-Indian Institute of Chemical Biology, Kolkata, India; Structural Biology and Bio-informatics division, CSIR-Indian Institute of Chemical Biology, Kolkata, India; Metabolic Disease Laboratory, Cell Biology and Physiology Division, CSIR-Indian Institute of Chemical Biology, Kolkata, India; Department of Molecular Biology, Netaji Subhas Chandra Bose Cancer Research Institute (NCRI) Kolkata, India

**Author notes:** These authors contributed equally (Diptankar Bandyopadhyay, Sudarshana Basu).

**Keywords:** Dicer1, Extracellular export, Ago2 interaction with Dicer1, Lipid Droplet, miRNA activity modulation, Phase separation

## Abstract

miR-122 is a liver specific miRNA that plays an important role in controlling metabolic homeostasis in mammalian liver cells. Interestingly, miR-122 on exposure to lipotoxic stress is reduced in liver cells. To fight stress, miRNA processor Dicer1 is depleted to cause reduced miR-122 production and the lowering of miRNA level ensures a better stress response in hepatocytes under lipotoxic stress. Interestingly, lipid droplets, formed in the liver cells on exposure to high fat, ensure cytoplasmic phase separation of Ago2 and prevent interaction of Ago2 with Dicer1. Lipid droplets bind miRNA and enhance miRNA-Ago2 uncoupling and Ago2 phase separation. Loss of interaction between Ago2 and Dicer1 eventually facilitates export and lowering of cellular Dicer1, a process also dependent on the endosomal maturation controller protein Alix, thereby ceasing pre-miRNA processing by Dicer1 in lipid exposed cells. Depletion of lipid droplets by downregulation of Perilipins with siRNAs resulted in a rescue of cellular Dicer1 level and Ago2-Dicer1 interaction. This is a novel mechanism that liver cells adopt to restrict cellular miRNA levels under stress condition. Thus, lipid droplets prevent cell death upon exposure to high fat by reducing intra and extracellular pool of miR-122 in hepatic tissue.

## Introduction

Lipid metabolism is centrally controlled in the liver where numerous lipid catabolic and anabolic processes as well as lipid export and import pathways function simultaneously to maintain lipid homeostasis in the human body. Deregulation of this balance leads to hepatic accumulation of lipids, a condition known as Non-Alcoholic Fatty Liver Disease (NAFLD). Non-Alcoholic Steatohepatitis or NASH is one form of NAFLD characterized by inflammation and hepatocyte ballooning. Development of NASH indicates increased risk for the further development of hepatic fibrosis, cirrhosis, liver failure and even hepatocellular cancer.

miRNAs are 20-22 nt long non-coding regulatory RNAs that fine-tune gene expression by binding to the 3’-UTR of target mRNAs and cause their repression or degradation (Bartel, 2018). miRNA biogenesis is a tightly regulated process both at the transcriptional and post-transcriptional level and disruption of this regulation leads to various human pathologies (Bartel, 2009; Esquela-Kerscher and Slack, 2006; Filipowicz et al., 2008). Dicer1, a Ribonuclease III (RNase III), processes pre-miRNA and loads the mature miRNA onto Ago2 to form the functional miRNA induced silencing complex (miRISC) in the cytoplasmic compartment of human cells (Kim et al., 2009) and is known to be targeted for controlling cellular miRNA content (Ghosh et al., 2013). Majority of mammalian genes are under miRNA regulation (Bartel, 2009) and miRNAs are also known to control diverse processes in hepatic lipid metabolism including *de novo* lipogenesis, fatty acid uptake, fatty acid oxidation and triglyceride export (Liu et al., 2015).

miR-122, identified as a highly abundant liver-specific miRNA, has a significant role in controlling hepatic metabolic processes (Chang et al., 2004; Jopling, 2012). Inhibition of miR-122 leads to down-regulation of a number of lipogenic and cholesterol biosynthesis genes in the liver (Esau et al., 2006). This in turn leads to reduced plasma cholesterol, increased hepatic fatty-acid oxidation and reduced synthesis of hepatic lipids. The SREBP-miR33 regulatory axis is also a well studied regulatory loop that maintains fatty-acid and cholesterol homeostasis in liver (Rottiers and Naar, 2012). miR-21 has been identified as another key miRNA that was decreased in high-fat diet fed mice and restoration of miR-21 blocked the stearic acid induced lipid accumulation (Ahn et al., 2012). Several previous studies on obesity-related NAFLD and NASH patients or on mouse models reveal alterations in the expression of various miRNAs in diseased livers (Cheung et al., 2008).

Lipid droplets (LDs) are highly conserved organelles originating from ER membranes that are comprised of a neutral lipid core formed by triacylglycerols and sterol esters and a surrounding monolayer of phospholipids (Martin and Parton, 2006; Wilfling et al., 2014). LDs prevent lipotoxicity in cells by sequestering excess free fatty acids as triacylglycerol (Jarc et al., 2018; Listenberger et al., 2003). Defects in lipid storage in LDs can lead to diseases such as type II diabetes, cardiovascular diseases and NAFLD (Krahmer et al., 2013). Advancement in the field of LD biology has revealed its importance in areas beyond lipid storage. LDs are now thought to be highly dynamic organelles having contact sites with subcellular structures and other organelles and host a repertoire of protein factors which regulate various signalling and metabolic pathways (Welte, 2007). LDs also act as sequestration sites for inactivation of certain nuclear proteins such as histones (Li et al., 2012) and MLX family of transcription factors (Mejhert et al., 2020). However, significance of increased number of lipid droplets reported in lipid exposed hepatic cells in connection to alteration in miRNA-mediated post-transcriptional gene regulatory processes remains entirely unclear.

As lipid metabolism is known to be tightly regulated by miRNAs, lipid accumulation could, in reverse, influence miRNA biogenesis and function. But the possible mechanistic link remains underexplored. In this study, we have addressed the question of how lipid challenged Huh7 cells alleviate stress due to lipid accumulation by exporting out miR-122 to buffer the cellular miRNA levels and thereby balance the metabolic processes. We have also explored the effects of short term or long-term exposure of hepatocytes to lipotoxic stress in terms of possible alterations in miR-122 target gene network. In this regard, we have identified a new role of LDs in controlling cellular miRNA content. By ensuring a phase separation of Ago2, LDs cause Ago2-Dicer1 interaction loss resulting in impairment of miRNA Loading Complex (miRLC) assembly in lipid loaded hepatic cells. This ensures a loss of Dicer1 and subsequently a reduced miR-122 level in hepatic cells exposed to high lipid. In correlation, MCD diet fed mice also showed decreased cellular miR-122 and Dicer1 in liver together with increased miR-122 in circulation.

## Results

### Increased extracellular export and associated drop in cellular miR-122 content in lipid exposed liver cells

miR-122 controls cholesterol biogenesis and to explore the reverse effect of cholesterol on hepatic miRNA machineries, we have treated Huh7 cells with various concentrations (0x, 1x, 2x and 5x) of cholesterol-lipid concentrate for 4h to determine the level at which there could have been a change in miRNA content. Exogenous cholesterol is known to be taken up via receptor-mediated endocytosis or by bulk flow endocytosis (Strauss et al., 2010) and stored as cholesterol esters in globular structures called lipid droplets in liver cells. Quantitative estimation of various miRNAs by qRT-PCR in cholesterol exposed and control cells revealed decreased levels of miR-122 with increase in cholesterol-lipid concentration used for treatment (Figure 1A and C). Relative quantification by qRT-PCR indicated a decrease in mature miR-122 levels with an increase in the duration of cholesterol treatment also (Figure 1B). All miRNAs tested, except miR-33 and miR-16, show a similar trend of reduced cellular levels against increasing cholesterol concentration (Figure 1C). Free fatty acid treatment (treatment with BSA conjugated Palmitic acid) also decreases mature miR-122 in Huh7 cells in a dose-dependent manner (Figure 1E). Treatment of mouse primary hepatocytes with cholesterol or palmitate also led to a decrease in mature miR-122 level (Figure 1G-H). Other miRNAs also showed a decrease in their content upon cholesterol treatment of primary hepatocytes. Surprisingly, upon lipid treatment, the decrease in miR-122 was not accompanied by a corresponding decrease in its precursor level. qRT-PCR revealed that precursor miR-122 levels increased with exposure to exogenous cholesterol or palmitate (Figure 1D and F), even when there has been a decrease in the mature form of the miRNAs. This may be indicative of decreased processing of the precursor miR-122; thereby leading to a ‘piling up’ of the same in lipid exposed cells. However, accumulation of pre-miRNA observed in both cholesterol and palmitate treated Huh7 cells can be contributed by a reduction in Dicer1 protein, the micro-processing enzyme in the pre-miRNA processor complex.

**Figure 1.**
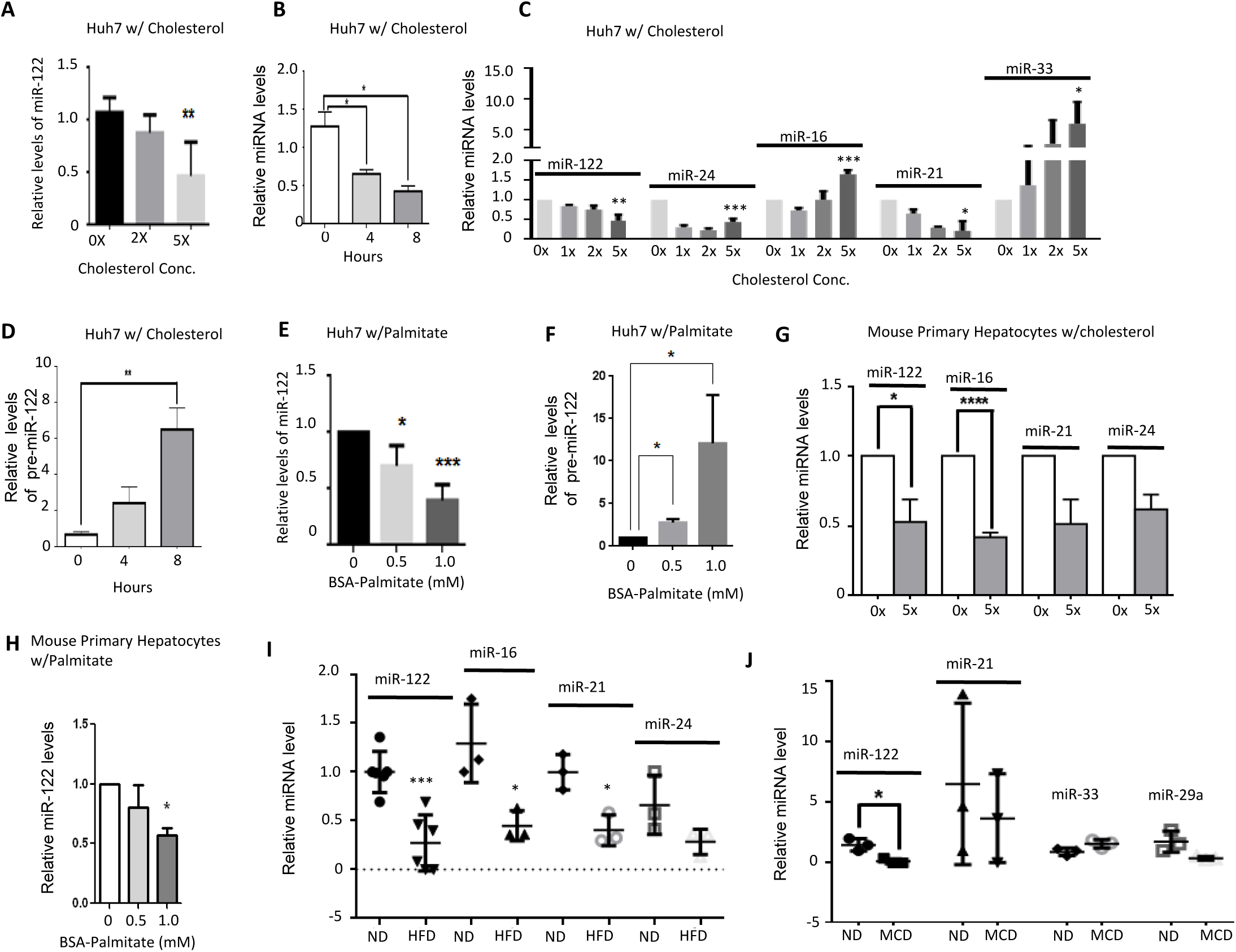
Drop of cellular miRNA upon lipid exposure in hepatocytes. **A** Cellular levels of miR-122 upon treatment of Huh7 cells with increasing concentration of cholesterol-lipid concentrate (0x, 2x, 5x) for 8 hours. miR-122 levels were quantified by quantitative RT-PCR detection from 25 ng of isolated cellular RNA (mean ± SD, n=4). Normalization was done against U6 snRNA. **B** Time dependent changes in cellular miR-122 content in 5x cholesterol-lipid concentrate treated Huh7 cells. miR-122 levels were quantified by quantitative RT-PCR detection from 25 ng of isolated cellular RNA (mean ± SD, n=4). Normalization was done against U6 snRNA. **C** Cellular levels of different miRNAs in Huh7 cells treated with cholesterol-lipid in a concentration dependent manner (0x, 1x, 2x, 5x) for 8 hours. All miRNA levels were quantified by qRT-PCR detection from 25 ng of isolated cellular RNA (mean ± SD, n=4). Normalization was done against U6 snRNA. **D** Precursor miR-122 cellular levels in Huh7 cells treated with 5x cholesterol-lipid concentrate for different time points. RT-qPCR detection of precursor levels was done from 200 ng of isolated cellular RNA (mean ± SD, n=4). 18S rRNA levels were used for normalization. **E** Concentration dependent changes in cellular miR-122 content in Huh7 cells treated with BSA conjugated palmitic acid (BSA-Palmitate) for 16 hours. For 0 mM treated Huh7 cells, equivalent amount of fatty acid free BSA was added. 25 ng of isolated cellular RNA was used for qRT-PCR measurement of miR-122 levels (mean ± SD, n=4). Normalization was done against U6 snRNA. **F** Cellular pre-miR-122 content in Huh7 cells treated with increasing concentration of BSA-Palmitate for 16 hours. Control Huh7 cells were treated with equivalent amount of fatty acid free BSA. Pre-miR-122 cellular levels were measured with 200 ng of cellular RNA by qRT-PCR and normalized against 18S rRNA levels (mean ± SD, n=3). **G** qRT-PCR based quantification of indicated miRNAs from mouse primary hepatocytes cultured *in vitro* and treated with or without 5x cholesterol concentrate for 8 hours. Cellular miRNA contents were quantified by qRT-PCR detection from 25 ng of isolated cellular RNA (mean ± SD, n=4). Data was normalized with respect to U6 snRNA. **H** Cellular miR-122 levels in mouse hepatocytes treated with either fatty acid free BSA (0 mM palmitate) or increasing concentration of BSA-Palmitate for 16 hours. qRT-PCR measurement of miR-122 was done with 25 ng of isolated cellular RNA (mean ± SD, n=4). Data normalized against U6 snRNA. **I** High Fat (HF) diet fed mice livers show reduced levels of miRNA. RT-qPCR detection of miRNAs was done from 25 ng of isolated cellular RNA. N=3 replicates each for chow (normal) diet fed mice and HF diet fed mice. Data represents Mean ± SD. Normalization was done by U6 snRNA. **J** Lipid accumulation in the liver of Methionine Choline deficient (MCD) diet fed mice leads to reduction in miRNA levels. RT-qPCR detection of miRNAs was done from 25 ng of isolated cellular RNA. N=3 replicates each for chow (normal) diet fed mice and MCD diet fed mice. Data represents Mean ± SD. Normalization was done by U6 snRNA For statistical significance, minimum three independent experiments were considered in each case unless otherwise mentioned and error bars are represented as mean ± S.D. P-values were calculated by utilising Student’s t-test. ns: non-significant, *P < 0.05, **P < 0.01, ***P < 0.0001.

### Cholesterol and Palmitiate treatment leads to exocytosis of DICER1 resulting in depleted cellular levels of the protein in mammalian hepatic cells

We hypothesized that the accumulation of pre-miR122 upon cholesterol treatment was probably due to reduced levels of Dicer1, a RNase III endonuclease responsible for cleaving pre-miRNA into mature miRNA. Intracellular levels of Dicer1 in cholesterol treated Huh7 cells showed a gradual decrease with increase in the time of exposure to cholesterol (Figure 2A). Both concentration and time dependent decrease in cellular Dicer1 levels on exposure to palmitate were also noted (Figure 2B-C). Similar decrease in Dicer1 levels was noted in primary hepatocytes exposed to increasing concentration of cholesterol or palmitate (Figure 2D). Extracellular export of Dicer1 could be an effective way of reducing its cellular content. To ascertain that the loss of cellular Dicer1 was due to its extracellular export, we measured the levels of Dicer1 in extracellular vesicles (EVs) isolated from culture supernatants of Huh7 cells grown in the presence or absence of cholesterol. We used HA-tagged Dicer1 transiently expressed in Huh7 cells to follow its export. Cholesterol treatment led to decreased levels of the HA-Dicer1 in Huh7 cells (Figure 2E). HA-Dicer1 levels were found to be higher in EVs isolated from cholesterol treated cells as opposed to EVs from control cells (Figure 2F). We also noted increase in EV associated HA-Dicer1 in Huh7 cells exposed to palmitate for 16 h (Figure 2G). Exposure of Huh7 cells to cholesterol or palmitate leads to extracellular export of miR-122 and increased miR-122 content has been documented in EVs isolated from palmitate or cholesterol treated Huh7 cells (Figure 2H).

**Figure 2.**
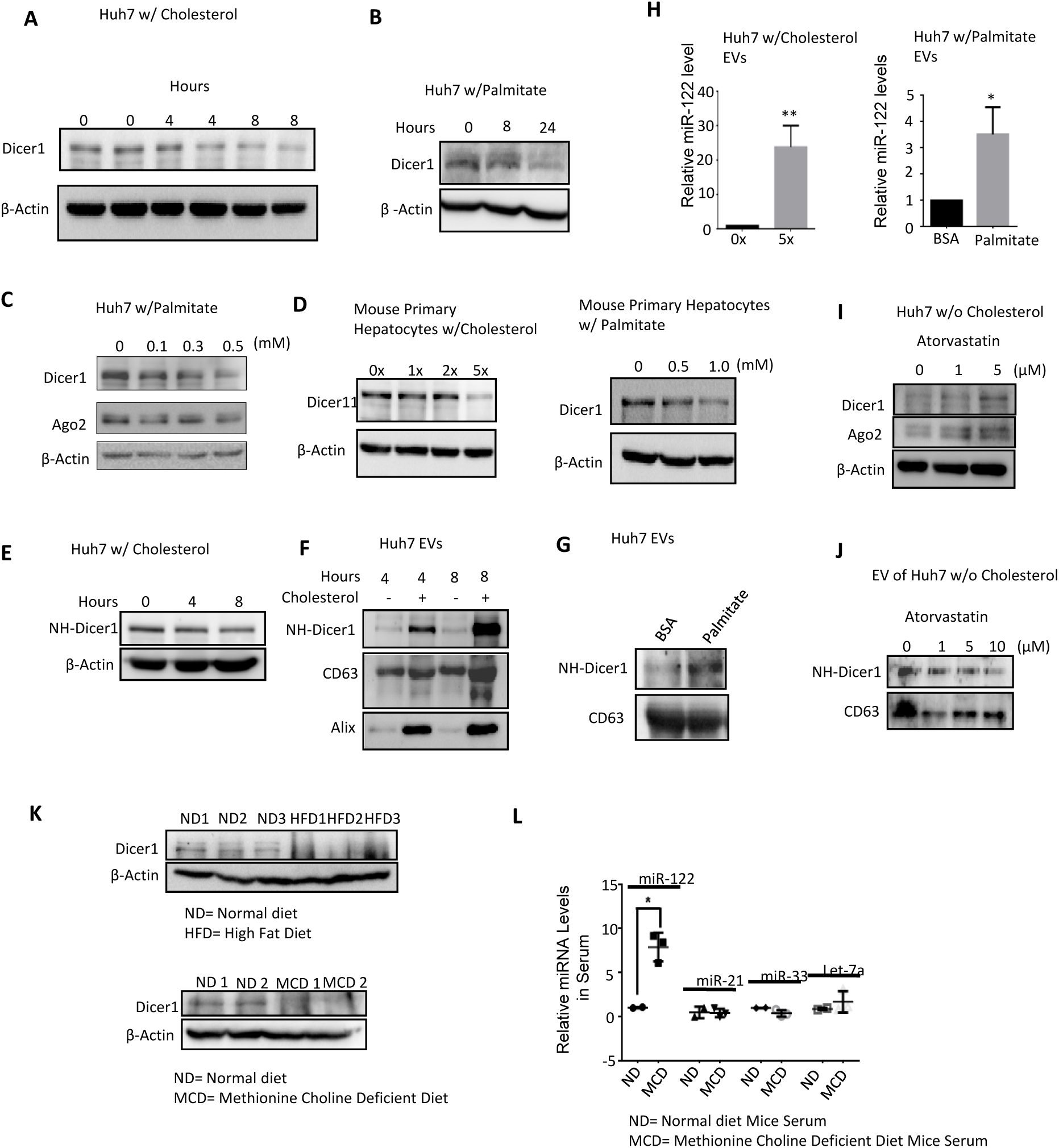
Reduction in cellular Dicer1 level in lipid exposed hepatocytes can account for reduced miR-122 level. **A** Western blot analysis of Dicer1 levels in control Huh7 cells and 5x cholesterol-lipid treated Huh7 cells for indicated time points. β-Actin was used as loading control. **B-C** Dicer1 levels in Huh7 cells treated with BSA-Palmitate in a time dependent (B) and dose dependent manner (C). Western blotting was done to measure the indicated protein levels. β-Actin was used as loading control. **D** Cellular Dicer1 levels on exposure to lipid L in mouse hepatocytes. Western blotting was done to estimate the amount of cellular Dicer1 levels of mouse primary hepatocytes treated with increasing concentration of cholesterol-lipid concentrate (left panel) or BSA-Palmitate (right panel). β-Actin was taken as loading control. **E-F** Cellular and extracellular protein levels of exogenously expressed NH-Dicer1 from Huh7 cells treated with cholesterol-lipid concentrate for indicated time points. Panel E shows western blot of cellular NH-Dicer1 levels in control and cholesterol-lipid treated Huh7 cells. Panel F shows EV associated NH-Dicer1 levels. Western blotting was done with anti-HA antibody to detect the amount of NH-Dicer1. β-Actin was used as loading control for cellular samples. CD63 and Alix were used as markers for EVs. **G** Effect of BSA-Palmitate treatment on NH-Dicer1 content of EVs. EVs from control (fatty-acid free BSA treated) and BSA-Palmitate treated Huh7 cells were isolated, protein was extracted and western blot analysis of NH-Dicer1 was done using anti-HA antibody. CD63 was used as a marker for isolated EVs. **H** EV associated miR-122 levels from control and cholesterol treated Huh7 cells (left panel) or control and BSA-Palmitate treated Huh7 cells (right panel). EVs were isolated from culture supernatant of Huh7 cells treated either with cholesterol or BSA-Palmitate, RNA was isolated and qRT-PCR detection of miR-122 was done from 100 ng of isolated RNA (mean ± SD, n=3). Data was normalized with respect to protein content of EVs isolated in each case. **I** Cellular Dicer1 and Ago2 levels of Huh7 cells treated with indicated concentration of Atorvastatin were detected by western blotting. β-Actin was used as loading control. **J** Western blot analysis of EV associated NH-Dicer1 levels from Huh7 cells treated with indicated concentration of Atorvastatin. CD63 was blotted as a marker of EV. **K** Dicer1 levels in Normal Diet (ND), High Fat (HF) diet and Methionine Choline deficient (MCD) diet fed mice liver were quantified. Western blot detection of Dicer1 was done from 50 μg total protein of mice liver lysates. **L** Quantification of circulating miRNAs in serum of MCD diet fed mice. Serum was isolated from chow and MCD diet fed mice. RT-qPCR detection of miRNAs was done from 100 ng of serum associated RNA. N=2 replicates for chow diet fed mice and 3 replicates for MCD diet fed mice. Data represents Mean ± SD. For statistical significance, minimum three independent experiments were considered in each case unless otherwise mentioned and error bars are represented as mean ± S.D. P-values were calculated by utilising Student’s t-test. ns: non-significant, *P < 0.05, **P < 0.01, ***P < 0.0001.

To verify the fact that excess lipid load induces Dicer1 export from hepatocytes, we performed experiments in presence of Atorvastatin, a known chemical inhibitor of HMGCR- the enzyme that catalyzes the rate-limiting step of cholesterol biosynthesis. Atorvastatin treatment is thus linked to reduced intracellular cholesterol level. Concentration dependent treatment of Atorvastatin to Huh7 cells in serum free medium led to an accumulation of cellular Dicer1 (Figure 2I). Concomitantly, there was a lowering of NH-Dicer1 in EVs of Atorvastatin treated Huh7 cell (Figure 2J). Nanoparticle Tracking Analysis (NTA) of EVs isolated from culture supernatants of control and lipid treated Huh7 cells revealed that lipid exposure does not alter the size distribution or concentration of EVs released by both treated or control cells (Supplementary Figure 1A-B). Overall, our results suggest increased export of Dicer1 that account for decreased cellular levels of the protein, which in turn reduces cellular miR-122 content in lipid challenged hepatocytes.

that in turn reduces cellular miRNA content in lipid challenged hepatocytes.

### Hepatic Dicer1 and miR-122 loss in the mice models of NAFLD

It seems that high lipid content has a negative effect on cellular Dicer1 level in hepatic cells. Does similar decrease of liver Dicer1 occur in mice exposed to high fat? We sought to examine the effect of elevated dietary lipids on hepatic Dicer1 in a high fat diet (HFD)- induced fatty liver mouse model. High fat diet fed mice liver exhibited reduced levels of Dicer1 (Figure 1K). This was accompanied by a significant lowering of various hepatic miRNAs (Figure 1I). Reduced levels of hepatic miRNAs reflects a long-term decrease in miRNA biogenesis, probably stemming from the fact that Dicer1 levels are lowered in the tissue. The most widely used diet to induce NAFLD/NASH in a rapid time is the methionine-choline deficient (MCD) diet. Hepatic lysates from MCD diet fed and normal chow diet (control) fed mice were western blotted to detect levels of Dicer1. Similar to what observed in high fat diet mice liver, a depleted level of the protein Dicer1 was observed in the liver of MCD diet mice as compared to control (Figure 1K). Levels of miR-122 and miR-29a were also found to be reduced in the corresponding liver tissue (Figure 1J). Serum levels of miR-122 were found to be significantly elevated in the disease model (Figure 2L). RT-qPCR analysis was done to detect changes in the levels of various lipid metabolic genes known to be altered with NAFLD. Elevated levels of the cholesterol biosynthetic enzyme HMGCo-A Reduactase (HMGCR) and the enzyme playing a key role in fatty acid transport across the inner mitochondrial membrane-Carnitine Palmitoyl Transferase (CPT1A) were observed (Figure 3A). Reduced levels of the enzyme catalyzing the rate-limiting step in the formation of monounsaturated fatty acids -Stearoyl Co-A Desaturase (SCD)- probably adds to hepatic stress due to high levels of saturated fatty acids (Figure 3A). Cholesterol 7 alpha-hydroxylase (CYP7A1) converts cholesterol to 7-alpha-hydroxycholesterol, the first and rate limiting step in bile acid synthesis. Levels of CYP7A1 were also reduced in the liver of mice fed with a methionine-choline deficient diet (Figure 3A). Hence the deregulation of lipid homeostasis is a natural consequence of hepatic lipid accumulation and probably arising from altered miRNA biogenesis induced by Dicer1 depletion in fat-loaded mouse livers.

**Figure 3.**
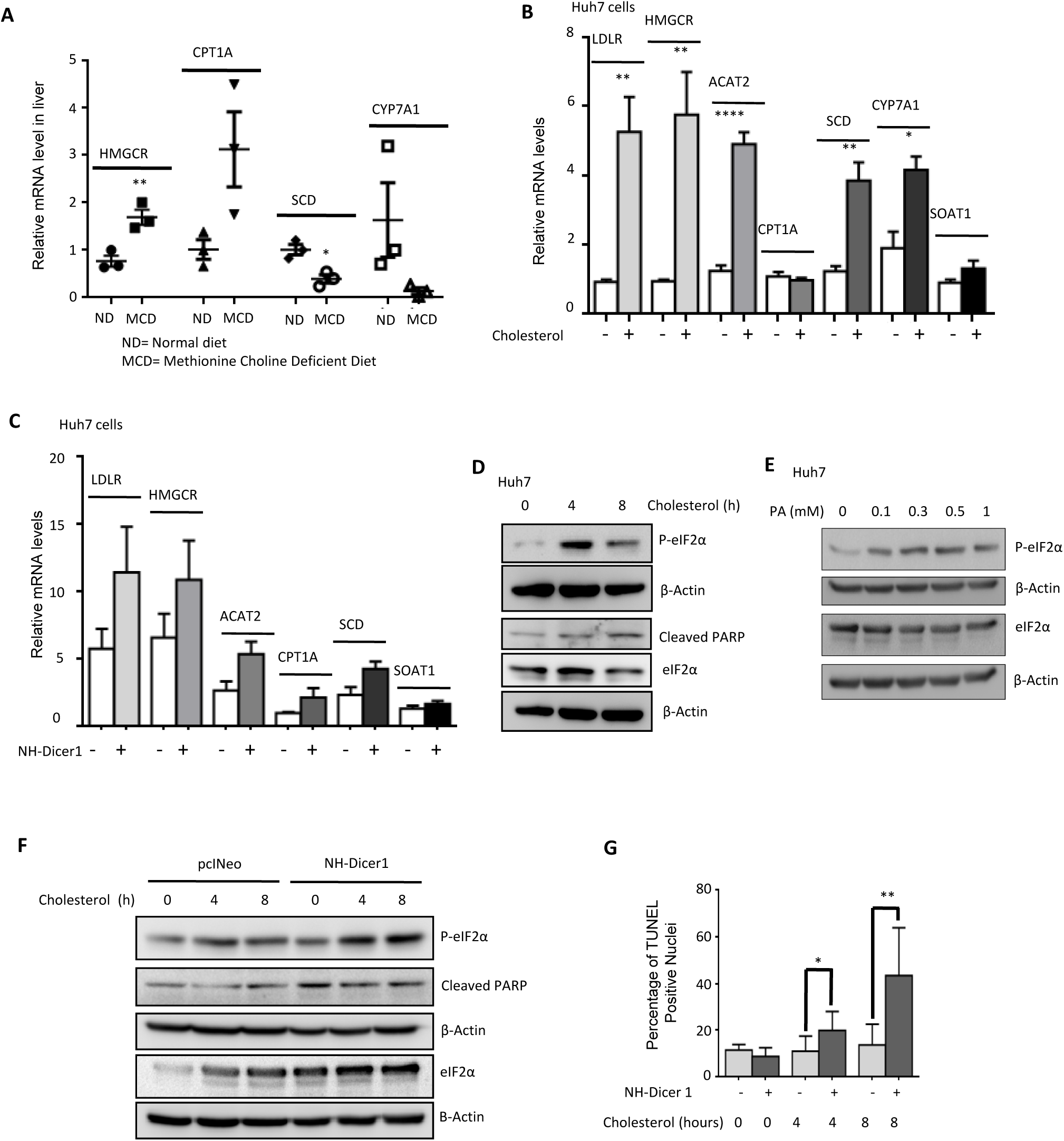
Exocytosis of Dicer1 counters ER stress in high lipid exposed hepatic cells. **A** Quantification of mRNAs of lipid metabolic genes in the liver of MCD diet fed mice. RT-qPCR detection using gene specific primers was done from 200 ng of RNA isolated from mice liver lysates. N= 3 replicates each for chow diet fed mice and MCD diet fed mice. Data represents Mean ± SD. Normalization was done using β-Actin mRNA. **B** Alterations in lipid metabolic genes induced upon treatment with 5x cholesterol-lipid concentrate for 4 hours. RT-qPCR detection of mRNAs using gene specific primers was done from 200ng of total cellular RNA. N= 5 replicates. Data shown represent Mean ± SD. Normalization was done using β-Actin mRNA. Abbreviations: LDLR; Low Density Lipoprotein Receptor, HMGCR; 3-Hydroxy 3-Methylglutary-CoA Reductase, ACAT2; Acetyl Co-A acetyltransferase 2, CPT1A; Carnitine palmitoyl transferase 1A, SCD; Stearoyl Co-A Desaturase, CYP7A1; Cholesterol 7 alpha-hydroxylase, SOAT1; Sterol-O-Acyltransferase 1. **C** Ectopic expression of NH-Dicer1 in lipid treated cells leads to amplified expression of deregulated lipid metabolic genes. Huh7 cells treated with 5x cholesterol-lipid concentrate were transfected with (1µg/ 1 x 10^6^ cells) plasmid encoding either NH-Dicer1 or control vector (pCI-Neo). RT-qPCR detection of mRNAs using gene specific primers was done from 200 ng of total cellular RNA. N= 5 replicates. Data shown represent Mean ± SD. Normalization was done using β-Actin mRNA. **D-E** ER stress and apoptosis is induced in cholesterol and palmitate treated Huh7 cells. Time dependent changes in EIF-2α, Phospho eIF2α (ER stress marker), and cleaved PARP (apoptosis marker) were detected by western blotting in 5X cholesterol-lipid concentrate treated Huh7 cells. β-Actin was used as loading control (D). Indication of ER stress was measured by western blot analysis of cellular levels of P-eIF2α and eIF2α from Huh7 cells treated with indicated concentrations of BSA-Palmitate (16 hours). For control Huh7 cells, equivalent amount of fatty acid free BSA was added. β-Actin was used as loading control (E). **F** NH-Dicer1 expression in lipid treated cells leads to elevated levels of the ER stress marker-phospho-eIF2α and apoptosis marker-cleaved PARP. Huh7 cells transfected with NH-Dicer1 expressing plasmid (1µg/ 1 x 10^6^ cells) or a control vector (pcINeo) were treated with 5X cholesterol-lipid concentrate for the indicated time points. Western blotting detection of respective proteins using specific antibodies was done. β-Actin was used as the loading control. **G** NH-Dicer1 expression in lipid treated cells lead to increased numbers of TUNEL positive nuclei. Huh7 cells were transfected with NH-Dicer1 expressing plasmid (1µg/ 1 x 10^6^ cells) prior to 5X lipid-concentrate treatment for indicated time points. Cells expressing HA-tags were detected by immunofluorescence using anti-HA primary antibodies and alexa-488 secondary antibodies (green). TUNEL staining was done and the percentage of green cells (NH-Dicer1 +ve) and non-green cells (NH-Dicer1 -ve) positive for TUNEL were identified. Graphical representation of the described experiment. N= 6 fields from 3 independent experiments. For statistical significance, minimum three independent experiments were considered in each case unless otherwise mentioned and error bars are represented as mean ± S.D. P-values were calculated by utilising Student’s t-test. ns: non-significant, *P < 0.05, **P < 0.01, ***P < 0.0001.

### Chronic lipotoxic stress may lead to changes in intra-cellular hepatic gene expression profile particularly the miR-122 target gene network

We have observed that on high cholesterol or fatty acid exposure, Dicer1 is exported and intra-hepatic levels of miR-122 tend to decline while its serum levels tend to increase. Such a re-distribution in the levels of miR-122, may result in large scale change in expression of its targets in the hepatic cells. Moreover, assuming that regulation of miR-122 is the key to control stress response in mammalian liver cells (Mukherjee et al., 2016), miR-122 lowering in hepatocytes should have commonalities in gene expression pattern with high fat exposed liver cells as high lipid exposure is known to be associated with cellular stress (Anderson et al., 2012; Fu et al., 2012). Thus, in order to get an overview regarding the changes in the miR-122 target gene expression in hepatic tissues under conditions of high fat exposure, differential gene expression analysis in the liver of mouse fed with high fat diet for 4 weeks, 12 weeks and 16 weeks (effect of cholesterol) was performed.

Short term or long term exposure of liver cells to high fat content resulted in up-regulation of 347, 2550, 184 and down-regulation of 143, 368, 38 genes due to high fat diet treatment for 4 weeks (HFD4), 12 weeks (HFD12) (GSE53381 and GSE93819 datasets) or high cholesterol (CH) exposure for 16 weeks (GSE58271), respectively. A higher number of genes were found to be commonly up-regulated compared to down-regulated genes (Supplementary Figure S2A). Since, high fat diet treatment has been found to be connected with down-regulated cellular levels of miR-122, genes found to be differentially expressed in hepatocytes upon high fat exposure were compared with probable miR-122 target genes (Wen and Friedman, 2012). Based on this analysis, we observed that nearly 29.37% [178] of the total possible up-regulated miR-122 target genes [606] show a significant change in their expression levels under high fat diet treatment conditions. Moreover, among the 256 genes that were found to be commonly up-regulated in two or more high fat diet or cholesterol treatment cases considered herein, 18.36% [47] of them were found to be miR-122 targets (Supplementary Figure S2A). Numbers of lipid metabolism related miR-122 target genes [7] were also found to be differentially regulated (Supplementary Figure 2A upper and lower panels). We found SCD2, a key regulator of energy metabolism and participating in fatty acid metabolism, to be up-regulated. Additionally, genes involved in downstream processing of cholesterol (CYP2B13; CYP2B9 and HSD17B6), ketone body catabolism (OXCT1) and arachidonic acid metabolism (CYP2B13; CYP2B9, CBR3) were also up-regulated. Fatty acid biosynthesis is likely to be down-regulated since ACACB (which regulates the rate-limiting step of fatty acid synthesis) is down-regulated (Supplementary Figure S2A lower panel). Further, pathway mapping of genes commonly up-regulated or down-regulated among these cases yielded additional insights into other de-regulated pathways possibly associated with down-regulated miR-122 levels upon high fat diet regime. Such de-regulated pathways involve broad categories like carbohydrate metabolism, cellular processes and signalling pathways (Supplementary Figure S2B). It was observed that the down-regulated miR-122 target genes are involved in carbohydrate and lipid metabolism regulation whereas up-regulated genes were found to be involved in cellular signalling or cellular process pathways (Supplementary Figure S2B). In particular, upon long term lipotoxic stress multiple miR-122 target genes involved in cellular pathways associated with different biological processes like cell motility, proliferation, differentiation, regulation of gene expression and survival like focal adhesion, tight junction or actin cytoskeleton regulation get de-regulated (Appendix Figure 1). Further, we observed that upon short term lipotoxic stress metabolic process related genes such as ACSS2, which regulates precursor (acetyl-coA) for fatty acid and cholesterol biosynthesis, gets down-regulated (Supplementary Figure S2B). Additionally, long term lipotoxic stress lead to de-regulation in miR-122 target genes like ACACB, UGT2B38 (down-regulated) and JUN, SPP1, COL6A3, PDGFRB, COL1A2, CTNNB1, CLDN2, CCND1, CCND2, PDGFRA, PRKCB, MAP3K1, CLDN7, COL1A1, ADCY7, CYBB, TLR2, NTRK2 (up-regulated) that could be important signalling cross-talk proteins participating in multiple pathways (Supplementary Figure S2B, Appendix Figure 2).

To verify the above trends observed in the differential expression analysis study, we evaluated mRNA levels of a few lipid metabolic genes in Huh7 cells treated with cholesterol-lipid concentrate for 4h (Figure 3B). mRNA levels of Low Density Lipoprotein Receptor (LDLR), 3-Hydroxy 3-Methylglutary-CoA Reductase (HMGCR), AcetylCo-A acetyltransferase 2 (ACAT2), Stearoyl Co-A Desaturase (SCD) and Cholesterol 7 alpha-hydroxylase (CYP7A1) were significantly increased (Figure 3B). LDL Receptors bind to and are responsible for the cellular intake of LDLs- the primary lipid carriers in circulation. HMGCR catalyzes the rate limiting step of cholesterol biosynthesis. Elevation of these proteins indicates increased lipid biosynthesis and uptake which serves to further increase the lipid load on the cell. ACAT2 is a cellular enzyme which converts cholesterol and fatty acids to cholesteryl esters. This is necessary to ameliorate the increased toxicity induced by the accumulation of cytosolic free lipids by esterifying the lipid for storage into lipid droplets. SCD catalyzes the rate-limiting step in the synthesis of unsaturated fatty acids. SCD deficiency has been reported to result in reduced adiposity, increased insulin sensitivity and resistance to diet-induced obesity (Flowers and Ntambi, 2008). Increase in SCD in the *in vitro* model system indicates increased lipid load on the cell. This increased lipid load is accompanied by slightly decreased fatty acid oxidation as indicated by the decrease in CPT1A- the enzyme responsible for ensuring the mitochondrial entry of fatty acids, the site of β-oxidation (Figure 3B). CYP7A1 converts cholesterol to 7-alpha-hydroxycholesterol which is the first and rate limiting step in bile acid synthesis. The synthesis of bile serves to reduce cholesterol level in hepatocytes. Sterol-O-Acyltransferase 1 (SOAT1) catalyzes the formation of fatty acid-cholesterol esters for subsequent storage into lipid droplets and thereby, regulates the equilibrium between free cholesterol and cytoplasmic cholesteryl esters. Increase in expression levels of these two genes (CYP7A1 and SOAT1) appear to be a physiological consequence of the elevated lipid content of the cell.

### Lowering of DICER1 acts as a cellular survival mechanism to counter the growing ER stress levels in lipid exposed cells

Our results demonstrate that increase in cellular lipid content leads to the export of Dicer1 from hepatocytes. But the possible role of this efflux and its physiological significance remained unknown. In an effort to answer this question, we overexpressed NH-Dicer1 in cholesterol-lipid treated cells and evaluated the mRNA levels of genes that have shown upregulation in high lipid conditions (Figure 3C). Overexpression of Dicer1 would serve to restore Dicer1 protein levels in lipid treated cells. However, contrary to our expectation, we found further elevations in levels of LDLR, HMGCR, ACAT2 and SCD in NH-Dicer1 expressing lipid treated cells (Figure 3C). This could have been possible only if the loss of Dicer1 was a pre-requisite to combat cellular stress. By overexpressing Dicer1, the stress condition may have been further increased as signified by elevated levels of lipid metabolic genes that were upregulated in lipid challenged hepatic cells.

High cellular cholesterol levels coupled with ER stress is being identified as a major cause behind the pathogenesis of various metabolic disorders (Sozen and Ozer, 2017). ER-stress-dependent dysregulation of lipid metabolism may lead to dyslipidemia, insulin resistance, cardiovascular disease, type 2 diabetes, and obesity (Basseri and Austin, 2012). It has been reported that the loading of cholesterol in macrophages induces the Unfolded Protein Response (UPR) due to the depletion of ER calcium stores (Feng et al., 2003). We wanted to determine if lipid loading in Huh7 cells leads to increased ER stress. To test that, we detected levels of phosphorylated eIF2α in cholesterol treated Huh7 cells. We observed increased phosphorylation of eIF2α along with elevated levels of cleaved PARP (Figure 3D). Palmitate treatment also resulted in the induction of ER stress (Figure 3E).

Therefore, lipid treatment appeared to lead to an increase in ER stress levels and apoptosis as detected in western blots of Phospho-eIF2α and cleaved PARP, respectively in liver cells (Figure 3D and E). Interestingly, expression of NH-Dicer1 was found to augment this stress response. Elevated levels of Phospho-eIF2α and cleaved PARP were detected in NH-Dicer1 transfected cholesterol treated Huh7 cells compared to cholesterol treated Huh7 cells that were transfected with a control vector (Figure 3F). TUNEL staining revealed increased number of TUNEL positive nuclei in Dicer1 overexpressed cells treated with cholesterol-lipid concentrate for 4 and 8 hours (Figure 3G). Cells transfected with HA-Dicer1 expression plasmids showed an increased percentage of TUNEL positive nuclei with respect to non-transfected cells upon lipid treatment (Figure 3G). Thus, extracellular secretion of Dicer1 acts as a survival mechanism adopted to counteract the growing lipotoxicity within the hepatic cell.

### Lipids reduce association of Dicer1 with the protein synthesizing Endoplasmic Reticulum

Exploring the mechanistic link between high lipid exposure and export of Dicer1 in hepatic cells, we did subcellular fractionation to check the cellular Dicer1 distribution in mammalian hepatic cells before and after lipid treatment. Surface of the rough endoplasmic reticulum (rER) has been reported as the site of miRNP nucleation and *de novo* miRNP formation (Bose et al., 2020; Stalder et al., 2013). rER also serves as the site where mRNAs interact with the repressive miRNPs before they are trafficked to endosomal compartment for subsequent degradation (Bose et al., 2017). Huh7 cells were used to prepare the microsome, a fraction enriched for rough endoplasmic reticulum. Microsomes have been identified as the sites for miRNA-mediated repression of target mRNAs and also the site for miRNP biogenesis (Barman and Bhattacharyya, 2015; Bose et al., 2017). We analysed the association of Dicer1 with microsomal fraction in presence and absence of cholesterol and palmitate and documented specific loss of Dicer1 from microsomal fraction in treated cells whereas no major change in Ago2 association with microsome was noted (Supplementary Figure S3A-B). These results suggest possibility of movement of Dicer1 from ER for its export. RT-qPCR analysis revealed higher Ct values for ER-associated miR-122, miR-16 and let-7a in palmitate treated cells (Supplementary Figure S3C). This indicates lower miRNA levels in the ER, probably due to their increased loading into EVs and less biogenesis in Palmitate treated cells due to Dicer1 loss. In subsequent analyses, isotonic cell lysates of Huh7 cells transiently expressing NH-Dicer1 were fractionated on a 3-30% iodixanol (Opti-prep^TM^) density gradient to separate out individual organelles. NH-Dicer1 was found to be mainly associated with the ER/ polysome fractions (based on colocalization with the ER protein Calnexin) in the control cell lysate (without cholesterol treatment). However, in the presence of cholesterol, this association was found to be reduced. Cellular NH-Dicer1 in cholesterol treated cell lysates was mainly localized in the endosomal/MVB fractions enriched for the marker protein Alix (Supplementary Figure S3D). Treatment of Huh7 cells with 0.5 mM palmitate also induced a similar shift of NH-Dicer1 from the ER to the endosomal/ MVB enriched fractions. Ago2 also showed a shifting towards the lighter fractions but its association with the ER remained unchanged between palmitate treated and non-treated cells (unpublished data).

To detect the endosomal association of Dicer1, Huh7 cell lysates (transiently expressing NH-Dicer1) were further fractionated on a 3-15% iodixanol (Opti-prep ^TM^) density gradient (Supplementary Figure S3E). The aim was to separate the early endosomal fractions from the late endosomal ones and determine the differential association of NH-Dicer1 with these fractions in presence of cholesterol. Fractions 2-4 were enriched for early endosomes (EE) (on the basis of colocalization with the marker protein Alix), and fractions 6-9 represented late endosomes (LE) positive for the marker Rab7A (Supplementary Figure S3E). We observed Dicer1 association with both fractions in the absence of cholesterol. However, in cholesterol loaded cells NH-Dicer1 was found to be predominantly lost from fractions 6 and 7 that are heavier than the EE positive fractions (Supplementary Figure S3E).

We performed microscopic analysis to visualize the spatial distribution of Endoplasmic Reticulum and endosomal compartments in lipid loaded cells to further explore the altered distribution of Dicer1 in lipid exposed hepatic cells. Huh7 cells expressing ER-DsRed and Endo-GFP were treated with cholesterol-lipid concentrate. Treated cells appeared to show a more peripheral endosomal distribution (green) with respect to control non-treated cells. In control cells, endosomes appeared as discrete organelles (green) associated with the ER (red) and homogenously distributed within the cell. However, in cholesterol treated cells they appeared either as large perinuclear bodies or aligned with the periphery of the cell (Supplementary Figure S3F). There also appeared to be reduced association of endosomes with the ER. This peripheral distribution of the endosome is also evident in Palmitate treated cells (unpublished data). We have reconstituted surfaces from the 3D image of the Endoplasmic Reticulum and endosomes by taking and analyzing images taken at various Z-levels. The surface reconstructed 3D image clearly shows a peripheral distribution of endosomes in cholesterol loaded Huh7 cells with respect to control untreated cells (Supplementary Figure S3G). Thus, lipid treatment leads to a rearrangement of the endosomes towards the periphery of the cell. It may be reasoned that the decrease in Dicer1 upon cholesterol treatment is probably connected to this altered distribution. Dicer1 associates with the peripheral endosome (as evident in gradient analysis experiments Supplementary Figure S3D and E) before getting exocytosed out of the cell.

### Dicer1 export is dependent on the endosomal protein Alix

The endosomal distribution of Dicer1 in cholesterol treated cells differs from that of untreated cells. The exact molecular partners involved in this altered localization needed to be characterized. To address the issue, proteins involved in the maturation of endosomes were depleted by transfecting Huh7 cells with specific siRNAs against Rab5A, Rab7A, HRS or Alix. Rab5A localizes to early endosomes where it is involved in the recruitment of Rab7A and the maturation to late endosomes (Hunker et al., 2006). Rab7A has been localized to late endosomes and functions as a key regulator in endo-lysosomal trafficking. It governs early-to-late endosomal maturation and endosome-lysosome transport through different protein-protein interaction cascades (Feng et al., 1995). HRS is involved in the endosomal sorting of membrane proteins into multivesicular bodies and lysosomes. HRS mediates the initial recruitment of ESCRT-I to endosomes and, thereby, indirectly regulates multivesicular body formation (Colombo et al., 2013; Hurley and Hanson, 2010; Pons et al., 2008; Scoles et al., 2000). It is also involved in the sorting of ubiquitinated proteins to clathrin-coated micro-domains of early endosomes (Sundquist et al., 2004). Alix, together with the lipid microenvironment, has been proposed to play a role in the formation of vesicles within multivesicular endosomes (Piper and Katzmann, 2007).

Depletion of Rab5A and HRS inhibits the decrease of Dicer1 observed in siControl transfected cholesterol treated Huh7 cells (Figure 4A-B). However there appeared to be a decrease in overall Dicer1 levels in Rab5A depleted cells even in the absence of cholesterol (Figure 4A). HRS depleted cells showed an increase in Dicer1 levels even in untreated cells (Figure 4B). Decrease of Dicer1 upon cholesterol treatment appears to be Rab7A independent (Figure 4C). Depletion of Alix by siRNA however was able to reverse the decrease in Dicer1 observed in Sicontrol transfected cells upon cholesterol treatment (Figure 4D). This reversal in Dicer1 levels was accompanied by a similar reversal in cellular miR-122 levels (Figure 4E, upper panel). These results led us to the conclusion that Alix inhibition may reverse the extracellular export of Dicer1 and miRNAs from lipid treated cells. To verify this, we isolated EVs from Alix depleted cells either treated or not treated with cholesterol-lipid concentrate. Levels of exported NH-Dicer1 and miR-122 were then analysed by western blotting and RT-qPCR, respectively. Cells depleted of Alix secreted reduced levels of NH-Dicer1 and miR-122, upon cholesterol treatment (Figure 4F). We detected levels of various other miRNAs to see if their exocytosis was also dependent on Alix. EVs isolated from siAlix transfected cholesterol treated cells showed reduced levels of secreted miR-16, 21 and 24 with respect to siControl transfected cholesterol treated cells (Figure 4G). In subsequent experiments, we found that the extracellular secretion of NH-Dicer1 from palmitate treated cells was also dependent on Alix (data not shown).

**Figure 4.**
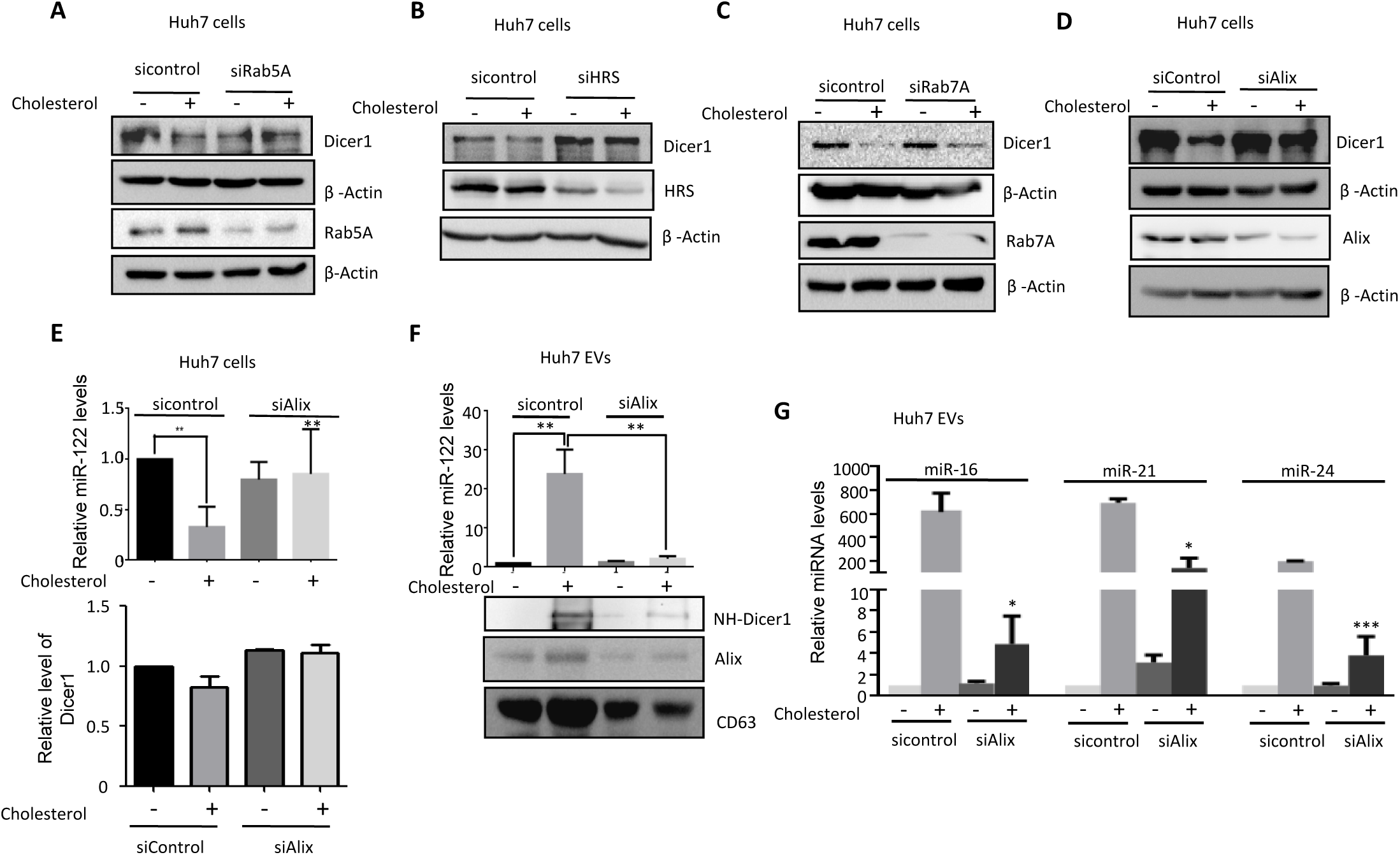
Cholesterol treatment of hepatic cells leads to increased extracellular release of Dicer1 through endosomal pathway. **A-D** Effect of inhibition of endosomal maturation on Dicer1 levels in control and 5x cholesterol-lipid treated Huh7 cells. Western blot analysis showing the cellular Dicer1 levels in control and 5x cholesterol-lipid treated Huh7 cells, depleted for Rab5A (A), HRS (B), Rab7A (C) or Alix (D). β-actin was used as loading control. Knockdown efficiency was also validated by western blot of the respective depleted factors. **E** Effect of siAlix treatment on cellular miR-122 and Dicer1 levels in control and 5x cholesterol-lipid treated Huh7 cells for 4 hours. Upper panel shows the relative miR-122 cellular levels quantified by qRT-PCR detection from 25 ng of isolated cellular RNA (mean ± SD, n=4). Normalization was done against U6 snRNA. Lower panel shows the quantification of relative Dicer1 levels from western blot analysis of cellular Dicer1 levels. β-Actin was used as loading control for normalization (data from multiple analysis). **F** Effect of siAlix treatment on EV content of miR-122 and NH-Dicer1 from control and 5x cholesterol-lipid treated Huh7 cells. EVs were isolated from culture supernatants of control and 5X cholesterol-lipid treated (4 hours) Huh7 cells transiently expressing NH-Dicer1, RNA was isolated and miR-122 levels were measured by qRT-PCR from 100 ng of isolated RNA and normalized against the protein content of isolated EVs (mean ± SD, n=4), upper panel. Western blot of EV associated NH-Dicer1 level is shown in lower panel. CD63 was used as a marker of isolated EVs. **G** EV content of different miRNAs (miR-16, miR-21, miR-24) from siControl and siAlix transfected Huh7 cells treated with or without 5X cholesterol-lipid for 4 hours. RT-qPCR detection of miRNAs was done from 100 ng of isolated RNA from EV extracts and normalized against the protein content of EVs. For statistical significance, minimum three independent experiments were considered in each case unless otherwise mentioned and error bars are represented as mean ± S.D. P-values were calculated by utilising Student’s t-test. ns: non-significant, *P < 0.05, **P < 0.01, ***P < 0.0001.

### Loss of Ago2-Dicer1 interaction leads to decreased Dicer1 processing and impaired loading of mature miR-122 to Ago2 to form functional miRNPs in lipid loaded Huh7 cells

In a way to reduce this export of miR-122, hepatic cells export out cellular Dicer1 to reduce cellular burden of this stress responsive miRNA to restrict tissue damage. In the process, mature miRNP levels drop and pre-miRNA levels increase. The key role of Dicer1, being an endoribonuclease, is to process the precursor miRNA duplex and load the mature miRNA strand onto Ago2 to form a functional miRNP complex (Bose and Bhattacharyya, 2016). Interaction of Dicer1 and Ago proteins has been studied extensively and has been found to be essential for miRNA function (Bose et al., 2020; Haase et al., 2005; Kolb et al., 2005). Surprisingly, we find that there is a substantial decrease in interaction between NH-Dicer1 and Ago2 in cholesterol-lipid treated and BSA-Palmitate treated Huh7 cells compared to their respective control Huh7 cells (Figure 5A and B). Ago2’s interaction with other partner proteins like P-body components largely remained unaltered upon lipid exposure (Figure 5C, left panel). We immunoprecipitated FA-Ago2 from control and cholesterol-lipid treated Huh7 cells and documented a reduction in the amount of miR-122 bound to FA-Ago2 in cholesterol-lipid treated Huh7 cells (Figure 5C, right panel). Thus, there might be a decrease in loading of miR-122 from Dicer1 to Ago2 due to their reduced interaction. Does loss of Ago2-Dicer1 interaction have a role in Dicer1 processing of pre-miR-122? Earlier, we observed an accumulation of pre-miR-122 in lipid treated Huh7 cells (Figure 1D and F). To delineate the mechanism, we immunoprecipiated NH-Dicer1 from cholesterol-lipid treated and control Huh7 cells and noticed an increase of miR-122 and miR-122* associated with NH-Dicer1 from cholesterol-lipid treated Huh7 cells (Figure 5D). This was consistent with a reduction of pre-miR-122 association with NH-Dicer1 in cholesterol-lipid treated Huh7 cells (Figure 5E). Since, Dicer1 is not able to load the mature miR-122 onto Ago2, probably due to reduced Ago2-Dicer1 interaction, so miR-122 remains associated with Dicer1 and as a result Dicer1 does not become available for the next round of processing of a new pre-miR-122 molecule.

**Figure 5.**
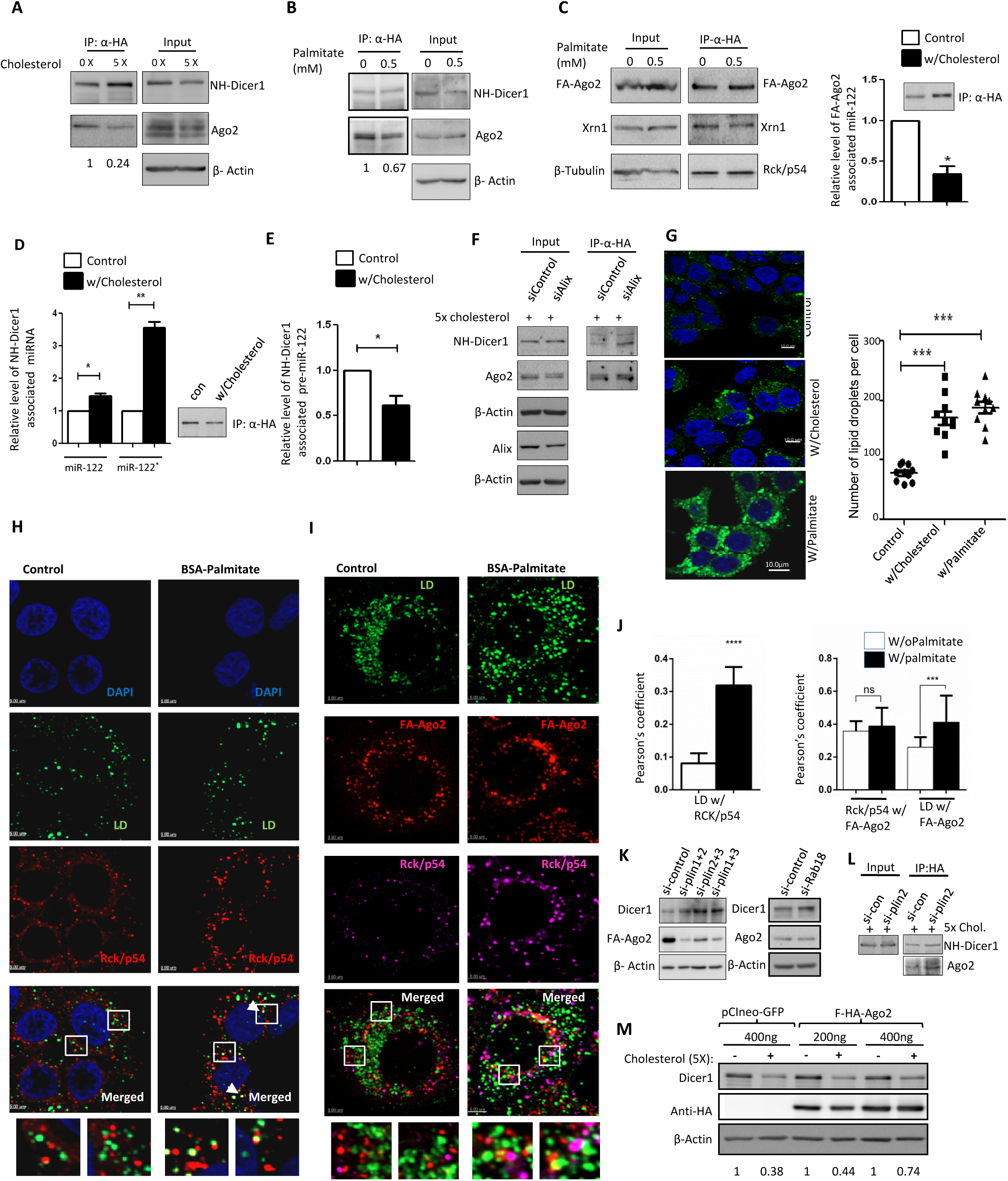
LD induced loss of interaction between NH-Dicer1 and Ago2 in lipid treated Huh7 cells causes a decrease in cellular Dicer1 level. **A-B** Effect of lipid treatment on the interaction between NH-Dicer1 and Ago2 proteins in Huh7 cells transiently expressing NH-Dicer1. Western blot analysis of immunoprecipitated NH-Dicer1 (using anti-HA antibody) and its associated Ago2 protein levels in control Huh7 cells and in Huh7 cells treated with either 5X cholesterol-lipid for 4hours (A) or BSA- Palmitate for 16 hours (B). Input levels of the indicated proteins were western blotted. Quantification of western blot bands was done and band intensity ratio of Ago2 is to NH-Dicer1 indicating amount of Ago2 co-immunoprecipitated per unit NH-Dicer1 was calculated. The values were normalized against the respective control sets and shown below the blots. **C** Interaction of transiently expressed FA-Ago2 with P-body proteins and miR-122 in control and lipid treated Huh7 cells studied by immunoprecipitation assay. Western blot analysis showing pulled down FA-Ago2 (using anti-HA antibody) and its associated P-body factors, Xrn1 and Rck/p54 in control and BSA-Palmitate treated Huh7 cells transiently expressing FA-Ago2. Input levels of the proteins are also shown (left panel). Right panel shows the effect of 5x cholesterol-lipid treatment on FA-Ago2 bound miR-122 in Huh7 cells transiently expressing FA-Ago2. Immunoprecipitation of FA-Ago2 (using anti-HA antibody) was done from control and 5x cholesterol treated (for 4 h) Huh7 cell lysates.. Immunoprecipitated samples were divided into two equal parts. Western blot was done with one part to quantify the amount of FA-Ago2 pulled down. RNA was isolated from the other part followed by quantification of miR-122 by qRT-PCR detection and normalization with respect to the amount of FA-Ago2 pulled down (mean ± SD, n=3). **D-E** Relative amount of miR-122, miR-122* and pre-miR-122 bound to NH-Dicer1 that were immunoprecipitated (using anti-HA antibody) from control and 5x cholesterol-lipid treated Huh7 cells expressing NH-Dicer1. Immunoprecipitated samples were divided into two equal parts. Western blot was done with one part to quantify the amount of NH-Dicer1 pulled down. miR-122, miR-122* (D) and pre-miR-122 (E) were quantified by qRT-PCR with the RNA isolated from other part and normalized to the amount of NH-Dicer1 immunoprecipitated (mean ± SD, n=3). **F** Western blot analysis of the amount of Ago2 co-immunoprecipitated with NH-Dicer1 in siControl and siAlix transfected Huh7 cells. Huh7 cells transiently expressing NH-Dicer1 were co-transfected with either siControl or SiAlix and the resulting cell lysates were immunoprecipitated with anti-HA antibody and western blotted for the indicated proteins. Input levels of NH-Dicer1, Ago2, Alix and β-Actin are also shown. **G** Bodipy 493/503 staining of lipid droplets (LDs) in Huh7 cells treated with either 5X cholesterol (4h) or 0.5 mM BSA-Palmitate (16h) or control cells (only BSA). Cells were counterstained with DAPI to visualize the nucleus. Fields represent one of 6 fields detected at 60x magnification. Scale bar represents 10µm. Quantification of number of LD has been done (n>100 cells) for all three conditions and plotted in the right panel. **H** Representative merged images depicting Rck/p54 protein localization with LDs in control and BSA-Palmitate (16h) treated Huh7 cells. LDs (green) were stained with Bodipy 493/503 and indirect immunofluorescence staining of endogenous Rck/p54 (red) was done. DAPI was used for staining the nucleus. Scale bar, 10µm. Marked areas are 5X zoomed. White arrows indicate regions of colocalization between LDs and Rck/p54 positive bodies. **I** Representative merged frames showing staining of LDs, Rck/p54 and FA-Ago2 in control and BSA-Palmitate treated (16h) Huh7 cells. LDs (green) stained with Bodipy 493/503 (green). Rck/p54 (magenta) and FA-Ago2 (red) were stained by indirect immunofluorescence of endogenous Rck/p54 and FA-Ago2, respectively. Scale bar represents 10 µm. **J** Pearson’s coefficient calculation showing mean ± SD values from at least three independent experiments (10 cells per experiment) of colocalization data between LDs and Rck/p54 (left panel) from experiment described in (H), FA-Ago2 and Rck/p54 (right panel) and FA-Ago2 and LDs (right panel) from experiment described in (I). **K** Levels of Dicer1, FA-Ago2 and endogenous Ago2 in Huh7 cells depleted of different LD associated factors. Left panel shows western blot analysis of Dicer1 and FA-Ago2 levels of Huh7 cells depleted of various combination of Perilipin isoforms (as indicated). Right panel shows levels of Dicer1 and Ago2, estimated by western blot of cell lysates from siControl and siRab18 transfected Huh7 cells. β-Actin was used as loading control. **L** Western blot analysis of immunoprecipitated NH-Dicer1 (using anti-HA antibody) and its associated Ago2 levels in cholesterol treated siControl and siPerilipin2 transfected Huh7 cells. Input levels of NH-Dicer1 were detected by western blotting. **M** Effect of ectopic expression of FA-Ago2 (200 and 400 ng for 0.5 x 10^6^ cells) or pCIneo- GFP (400 ng for 0.5 x 10^6^ cells) on endogenous Dicer1 level in Huh7 cells untreated or treated with 5x cholesterol-lipid concentrate for 4h. Relative levels of Dicer1 have been calculated from band intensities and normalized against β-Actin used as loading control. For statistical significance, minimum three independent experiments were considered in each case unless otherwise mentioned and error bars are represented as mean ± S.D. P-values were calculated by utilising Student’s t-test. ns: non-significant, *P < 0.05, **P < 0.01, ***P < 0.0001.

### Lipid Droplet association of Ago2 regulates cellular abundance of Dicer1

What causes the uncoupling of Dicer1 and Ago2? Why is Ago2 not available for interaction with Dicer1 and subsequent processing and loading of mature miRNA by Dicer1 to Ago2? There was no change in interaction between NH-Dicer1 and Ago2 in Alix depleted cholesterol treated Huh7 cells compared to siControl transfected cholesterol exposed Huh7 cells (Figure 5F). Therefore the Dicer1-Ago2 interaction loss must happen upstream of Dicer1 export, a process controlled by Alix. Treatment of Huh7 cells with lipid load in the form of cholesterol-lipid concentrate or BSA-Palmitate would lead to intracellular lipid accumulation. Upon staining of the cells with Bodipy^TM^ 493/503, a lipophilic dye which stains the neutral lipids present in the form of droplets, we found an increase in the number of these lipid droplets (LD) upon exposure of Huh7 cells to cholesterol-lipid concentrate or BSA-Palmitate (Figure 5G). Recent reports suggest that lipid droplets exhibit phase behaviour and liquid-crystalline phase transition of lipid droplets influence various cellular processes and its inter-organellar interactions (Mahamid et al., 2019). RNA processing bodies or P-bodies (PBs) are membraneless phase separated organelles that are known to harbor repressed mRNAs along with miRNPs and RNA degrading enzymes and take part in miRNA-mediated translation repression of target messages. In PBs, repressed mRNAs and Ago2 can be stored for future use (Bhattacharyya et al., 2006; Hubstenberger et al., 2017). Interestingly, we observed a strong interaction between LDs and PBs in Palmitate treated Huh7 cells (Figure 5H and J, left panel). Interaction between these two phase separated entities might aid in exchange of factors to influence each other’s biochemical roles. A previous study has shown the partitioning of Ago2 to lipid droplets is required for HCV replication in Huh7 cells (Berezhna et al., 2011). We demonstrate the selective increase in association of exogenously expressed FA-Ago2 with lipid droplets in palmitate treated Huh7 cells (Figure 5I and J, right panel). To check whether the lipid droplet play a part in modulating the abundance of Dicer1, we deplete lipid droplet by targeting perilipins, constituents of lipid droplets. Upon depleting Perilipins, the LD associated factor known to stabilize LDs (Kaushik and Cuervo, 2015; Kaushik and Cuervo, 2016), we found a rescue in cellular Dicer1 levels in cholesterol-treated Huh7 cells (Figure 5K left panel and Figure 7C). This is accompanied by an increase in Ago2 association of NH-Dicer1 in Perilipin2 depleted, cholesterol treated Huh7 cells (Figure 5L). Also, we detected an increase in cellular Dicer1 levels in Huh7 cells depleted for Rab18 –a protein known for its role in LD biogenesis (Figure 5K, right panel). Thus, increased interaction of Ago2 with LDs might cause reduced Ago2-Dicer1 interaction which in turn leads to increased Dicer1 export. To support this notion, we expressed FH-Ago2 and noted a partial restoration of cellular Dicer1 levels in cholesterol treated Huh7 cells (Figure 5M).

### Lipid droplets ensure phase separation of miRNA and Ago2 that leads to depletion of Dicer1 interacting cytoplasmic Ago2 pool

Owing to high lipid to protein ratio, LDs are of extremely low density and hence, LDs float on top of a sucrose density gradient upon ultracentrifugation. We isolated LDs by such a floatation gradient from previously published protocol (Ding et al., 2013) with minor modifications. We detected a significant amount of miR-122 that have been found to be enriched in LD fractions isolated from BSA-Palmitate treated Huh7 cells (Figure 6A-B). The absence of Ago2 in LD fraction could be attributed to the fact that Ago2 interacts transiently with LDs in a phase-interphase boundary which is lost during rigorous steps of biochemical isolation (Figure 6A). But how does miR-122 interact with LDs? RNase protection assay with LD fractions of palmitate stimulated Huh7 cells indicated the presence of miR-122 mostly on the surface of LDs and hence, are sensitive to RNase treatment (Figure 6C). Does Ago2-miRNA interaction also get altered in presence of LD? Or can LD help Ago2-miRNA uncoupling? We noted an overall decrease of miR-122 bound to FA-Ago2 immunoprecipitated from lipid exposed Huh7 cells (Figure 5C, right panel). Reduction of miR-122 associated with Ago2 in lipid loaded Huh7 cells might be a combination of reduced Dicer1 loading as well as active uncoupling of miR-122 from Ago2. To investigate the latter, we hypothesized that LDs might act as a miRNA adsorbing sponge and thus might be the site of miR-122 uncoupling from Ago2. We document a reduction in the level of miR-122 associated with immunoprecipitated FA-Ago2 upon increasing time of incubation of FA-Ago2 and miR-122 expressing HEK293 lysate with purified LDs from palmitate stimulated Huh7 cells (Figure 6D and E). Earlier, we documented a rescue of Ago2-Dicer1 interaction in Perilipin 2 depleted lipid exposed Huh7 cells. To assess the involvement of LDs to influence Ago2-Dicer1 interaction, we performed an *in vitro* Ago2-Dicer1 interaction assay. Incubation of HEK293 lysate expressing NH-Dicer1 with LDs isolated from palmitate stimulated Huh7 cells led to lesser amount of Ago2 co-immunoprecipitated with NH-Dicer1, compared to control reactions in absence of LDs (Figure 6F and G).

**Figure 6.**
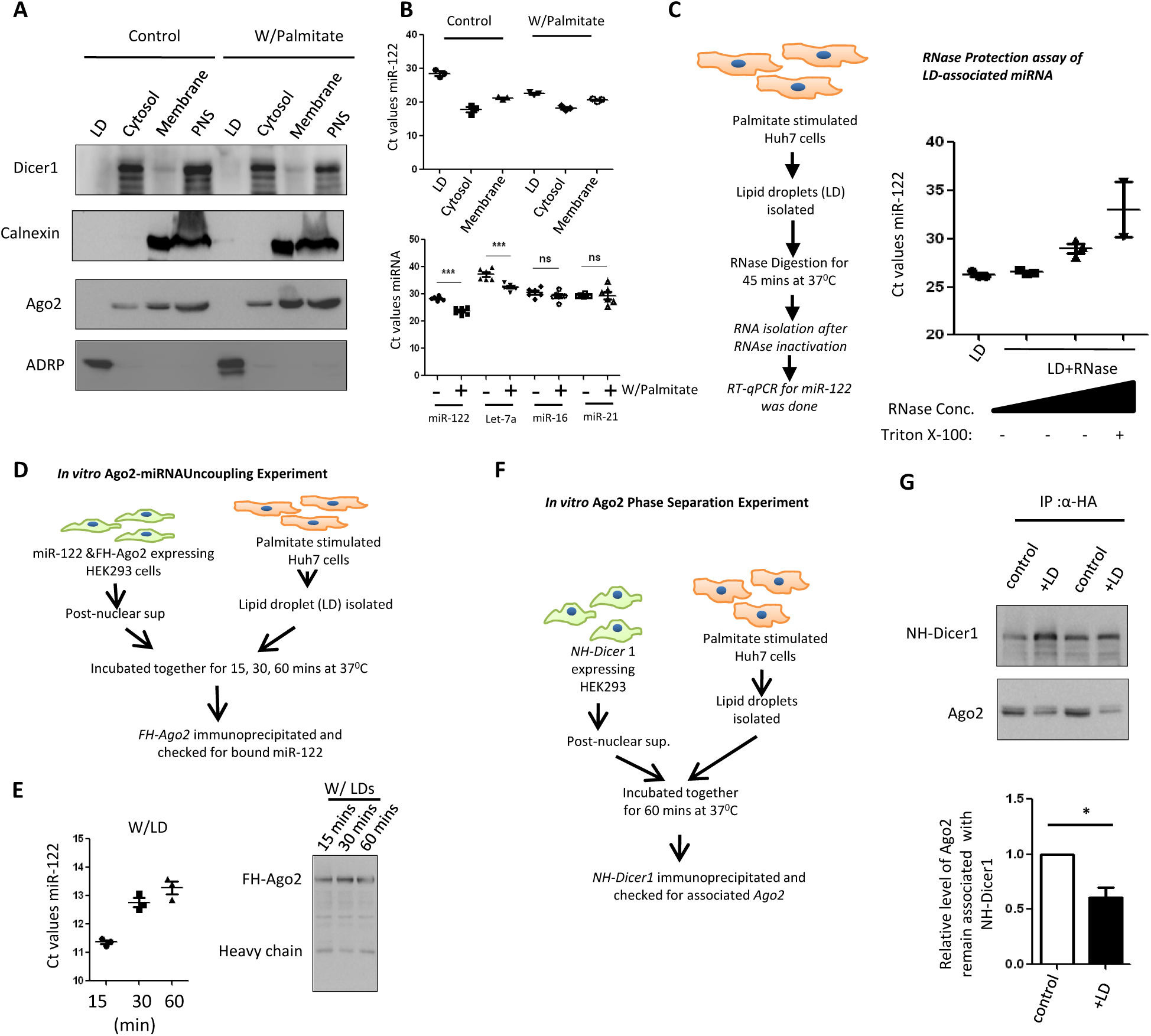
Lipid droplets prevent Ago2-Dicer1 interaction by phase separation of Ago2. **A** Western blot analysis to assess the purification of LDs from control and BSA-Palmitate treated (16h) Huh7 cells by following enrichment of adipose differentiation-related protein (ADRP)- a LD marker. Also, cytosolic, total membrane and post-nuclear supernatant (PNS) fractions are shown. Relative abundance of Dicer1 and Ago2 in the fractions was detected by western blots. ADRP and Calnexin were used as markers of LD and endoplasmic reticulum, respectively. **B** Endogenous miR-122 levels in the LDs, cytosolic, total membrane fractions were quantified by qRT-PCR from RNA isolated from equivalent amount of each fraction. The Ct values of miR-122 obtained were plotted (upper panel, mean ± SD, n=3). Similar Experiments were done for other miRNAs and related level change in LD associated miRNA content before and after palmitate treatment shown in the bottom panel (mean ± SD, n=3). **C** RNase protection assay of LD associated miR-122 done with LDs isolated from 16 hours BSA-Palmitate stimulated Huh7 cells. Scheme of the experiment is shown in left panel. Isolated LDs were digested for 45 mins at 37°C with RNase (10 µg or 20 µg/ml), RNase (20 µg/ml) in presence of Triton X-100 and no RNase control (LD), followed by RNA extraction. miR-122 levels in residual RNA were quantified by qRT-PCR and the Ct values obtained were plotted and shown in right panel (mean ± SD, n=3). **D-E** Time course experiment for demonstrating the effect of LDs on Ago2-miRNA uncoupling. Experimental scheme for the *in vitro* assay is shown in (D). miR-122 and FA-Ago2 expressing HEK293 post-nuclear supernatants (1×10^6^ cell equivalent amount in each reaction) were incubated with purified LDs (from BSA-Palmitate stimulated Huh7 cells, 5×10^6^ cell equivalent amount in each reaction) at 37° C for indicated time points. Post incubation, FA-Ago2 was immunoprecipitated and checked for bound miR-122 by qRT-PCR. Western blots show the amount of FA-Ago2 pulled down (E, right panel). Ct values of immunoprecipitated FA-Ago2 associated miR-122 are plotted (E, left panel) **F-G** *In vitro* Ago2 phase separation in presence of LDs alters its interaction with NH-Dicer1. Schematic depiction of the experiment is shown in F. Assay was performed at 37° C for 1h with NH-Dicer1 expressing HEK293 post-nuclear supernatant (1×10^6^ cell equivalent amount in each reaction) in absence or presence of LDs isolated from BSA-Palmitate stimulated Huh7 cells (5×10^6^ cell equivalent amount in each reaction) followed by immunoprecipitation with anti-HA antibody. Western blot of immunoprecipitated NH-Dicer1 and associated Ago2 is shown (G, top panel). Band intensity analysis from three independent experiments was done to calculate the amount of Ago2 that remained associated with NH-Dicer1 and plotted (G, bottom panel).

These data suggest that Ago2 gets phase separated by LDs and become less accessible for Dicer1 interaction. The phase separation of Ago2 by LD may cause a retardation of its dynamics and we have used single point Fluorescence Correlation Spectroscopy (FCS) analysis to study the the diffusion pattern and dynamics of transiently expressed GFP-Ago2 in control and palmitate treated Huh7 cells. The autocorrelation functions of GFP-Ago2, thus obtained, were fitted to an anomalous diffusion model, considering that the cytosol was crowded. The diffusion time through the confocal volume of GFP-Ago2 molecules in palmitate treated Huh7 cells was extremely slow compared to GFP-Ago2 molecules in control Huh7 cells. As a result, the diffusion coefficient (D) of GFP-Ago2 in palmitate treated Huh7 cells could not be calculated due to a retarded movement of GFP-Ago2 through the confocal volume resulting in a dispersed fit of autocorrelation curve in palmitate treated Huh7 cells (Supplementary Figure S4B and C). Increased interaction of GFP-Ago2 with a larger, slowly diffusing ensemble like LDs in palmitate treated Huh7 cells could be the reason for the slower diffusion of GFP-Ago2 through the confocal volume. To ascertain the involvement of LDs in phase separating out and modulating the diffusion pattern of GFP-Ago2 in palmitate exposed Huh7 cells, we performed single point FCS analysis in non-siRNA and siPerilipin 2 transfected, palmitate treated Huh7 cells. Interestingly, we found a recovery of diffusion (faster diffusion) of GFP-Ago2 molecules in Perilipin 2 depleted palmitate treated Huh7 cells (Supplementary Figure S4D) that is comparable to that of control Huh7 cells, indicated by the rescue of the diffusion coefficient (Supplementary Figure S4B). These results confirm the role of LDs in interacting with and phase separating out Ago2 in lipid loaded Huh7 cells.

### Elevated levels of HuR mediate exocytosis of Dicer1 in lipid loaded Huh7 cells

In Figure 6B we have measured the change in association of miRNas with LD isolated from control and palmitate treated hepatic cells. The human ELAV protein HuR is a well-known stress response factor which translocates to the cytoplasm to facilitate relief of repressed mRNAs in response to starvation (Bhattacharyya et al., 2006). It also bind miRNAs with specificity and has been reported to bind and ensure export of miR-122 and let-7a from hepatic Huh7 and non-hepatic HeLa cells (Mukherjee et al., 2016). Also, HuR is known to relocalize to the cytoplasm in case of lipid challenged cells (Lu et al., 2015). Do HuR plays a role in LD compartmentalization of miRNA facilitate Dicer1 export? We noted elevated cellular levels of HuR in Huh7 cells treated with either cholesterol-lipid concentrate or palmitate in a dose dependent manner (Figure 7A and B). To find out whether HuR has any role in modulating the abundance of Dicer1 in lipid treated cells, we performed experiments in HuR depleted Huh7 cells. Interestingly, depletion of HuR could inhibit the cellular decrease in Dicer1 levels in cholesterol-lipid treated Huh7 cells (Figure 7C). Ectopic expression of HA-HuR in Huh7 cells led to a decrease in cellular Dicer1 levels as well (Figure 7D). Previous reports suggest the importance of HuR in packaging and export of miR-122 from starved Huh7 cells (Mukherjee et al., 2016). To this end, we investigated the role of HA-HuR in export of Dicer1 from Huh7 cells. Surprisingly, ectopic expression of HA-HuR led to elevated levels of NH-Dicer1 in EVs compared to NH-Dicer1 content in EVs from pCIneo transfected Huh7 cells (Figure 7E). Also, HA-HuR expression led to a reduction in Ago2-Dicer1 interaction (Figure 7F), a phenomenon that was also observed in lipid treated Huh7 cells (Figure 5A and B).

**Figure 7.**
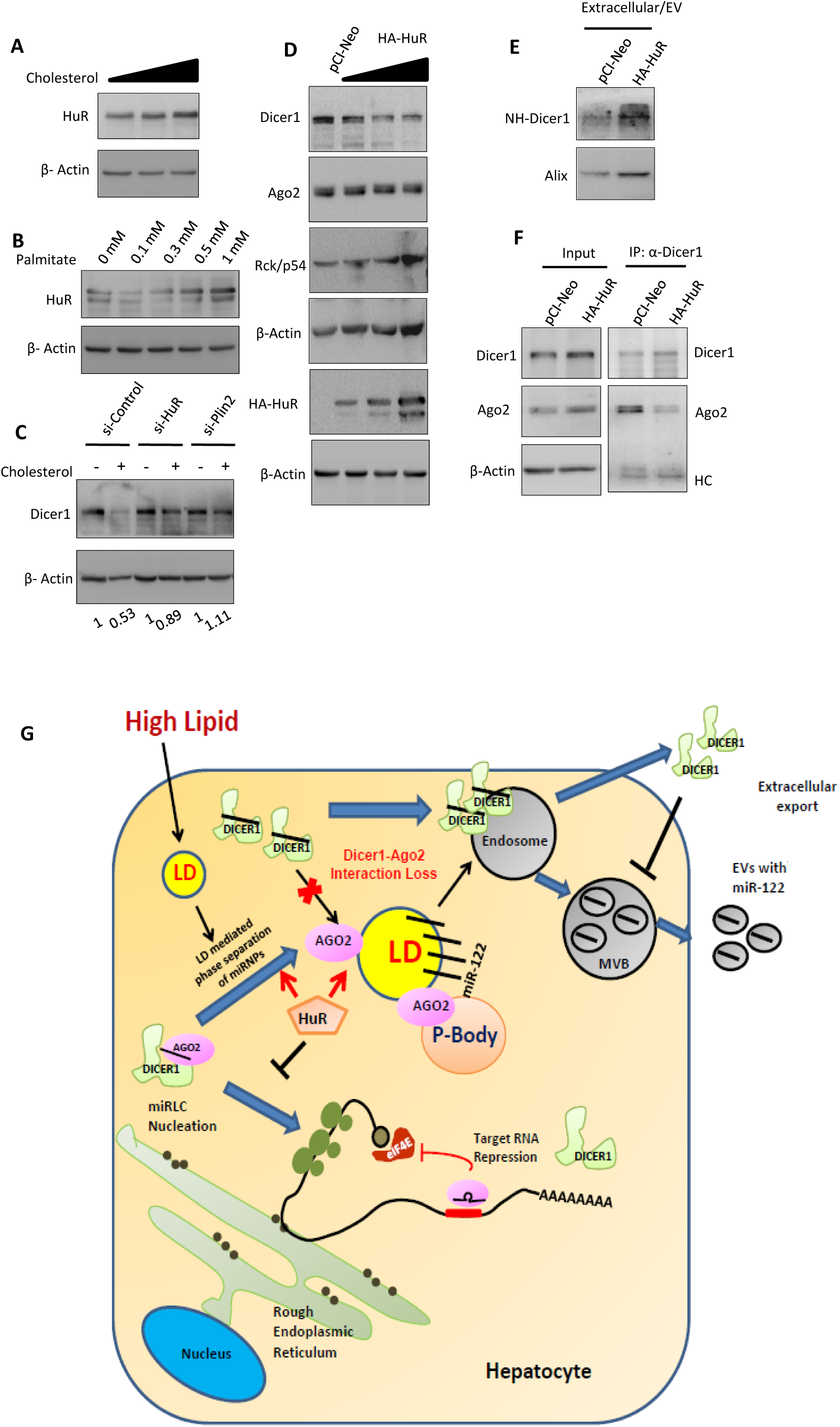
HuR mediated loss of Ago2 Dicer1 interaction in hepatic cells. **A-B** Western blot analysis of cellular levels of endogenous HuR protein in Huh7 cells treated with increasing concentration of cholesterol-lipid cholesterol for 4h (A) and BSA-Palmitate for 16h (B). β-Actin was used as loading control. **C** Cellular levels of Dicer1 protein estimated by western blot analysis from siControl, siHuR and siPlin2 transfected Huh7 cells treated with or without 5x cholesterol for 4h. β-Actin was used as loading control. Ratio of band intensities of Dicer1 is to β-Actin for each lane is given below the blot. **D** Western blot analysis showing cellular levels of Dicer1, Ago2 and Rck/p54 in Huh7 cells transfected with increasing amount of HA-HuR. pCIneo was transfected in control Huh7 cells. Expression levels of HA-HuR were confirmed by western blot using anti-HA antibody. β-Actin was used as loading control. **E** Effect of HA-HuR expression on EV content of NH-Dicer1. EVs were isolated from culture supernatant of Huh7 cells either expressing pCIneo or HA-HuR. Western blot analysis of EV associated NH-Dicer1 was done. Alix was used as an EV marker protein. **F** Effect of HA-HuR expression on the interaction between Ago2 and Dicer1 in Huh7 cells. Endogenous Dicer1 was immunoprecipitated from control and HA-HuR expressing cell extracts. Western blot analysis shows the amount of Ago2 co-immunoprecipitated with Dicer1 protein. Also shown are the input levels of the indicated proteins. pCIneo was transfected in control Huh7 cells. G A possible mechanism of how LDs accelerate the extracellular export of Dicer1 and thereby, reduces cellular miR-122 levels to minimise miR-122 export from lipid loaded hepatocyte. Excess lipid load in hepatocytes causes increased accumulation of LDs which ensure phase separation of Ago2. HuR mediated uncoupling of miRNAs from Ago2 promotes phase separation. Increased interaction of LDs with PBs may cause localisation of Ago2 to PBs These events prevents interaction of Dicer1 with Ago2, as a consequence of which Dicer1 translocates to the endosomes and gets exported out of the cells. The lowering of Dicer1 eventually lowers cellular miR-122 levels and restricts miR-122 mediated activation of neighbouring macrophages.

## Discussion

Findings described here have highlighted a unique mechanism adapted by hepatocytes to alleviate stress due to excess lipid load by lowering cellular miR-122 levels. The cells do so by actively exporting out miR-122 and the pre-miRNA processor Dicer1. Bioinformatic analysis revealed that miR-122 target genes which participate in metabolism associated processes (ACSS2, ACACB) were also down-regulated upon short term high fat exposure. However, miR-122 target genes (44) like CTNNB1, PDGFRA, PDGFRB, CCND1, PRKCB etc. involved in regulating cellular process or signalling pathways become up-regulated only upon prolonged exposure to high fat content. Thus, based on the computational analysis it appeared that the initial export of miR-122 from the hepatic cells might have protective effects by aiding in re-adjustment of cellular lipid levels. The later phase, the lowering of Dicer1 in lipid exposed hepatic cells, ensures the low-miRNA condition essential for stress response in hepatic cells (Figure 7G).

Modulation of Dicer1 availability, stability and abundance or tight regulation of Dicer1 activity is employed in cells to balance the mature miRNA expression. TAR RNA binding protein (TRBP) and protein kinase R-activating protein (PACT) are known to interact with Dicer1 and causes stabilization and enhanced processivity of Dicer1 (Haase et al., 2005) (Chendrimada et al., 2005). Various signalling pathways are also known to modulate the abundance and activity of Dicer1. Phosphorylation of TRBP by MAPK/ERK pathway is known to stabilize Dicer1 and increase Dicer1 activity leading to the expression of growth-promoting miRNAs and reduction of let-7a which is a tumour suppressor miRNA (Paroo et al., 2009). Variation in the 5’-UTR of Dicer1 mRNA and their restricted expression pattern in a tissue specific manner suggests another layer of regulation (Singh et al., 2005). The pluripotency factor Lin-28 is also known to inhibit processing of pre-let 7 RNA by Dicer1 in embryonic stem cells and embryocarcinoma cells (Rybak et al., 2008). In another report it has been shown how a protozoan parasite specific protease reduces hepatic cell Dicer1 to inhibit biogenesis of miR-122 and thereby, stop cholesterol production in liver cell. This is important for sustained infection by the pathogen where restoration of Dicer1 in infected liver can clear the infection (Ghosh et al., 2013). The present study reports extracellular export of Dicer1 and miR-122 as a novel way of buffering cellular miRNA levels in lipid loaded Huh7 cells. While miRNAs are known to regulate lipid metabolic processes, our findings suggest the existence of a reciprocal mechanism of controlling miRNA levels in lipid exposed hepatic cells.

miRNA loading complex (miRLC) comprising of Dicer, TRBP and Ago2 can process a pre-miRNA and load mature miRNA onto Ago proteins to cause target RNA cleavage in an ATP-independent manner when exposed to pre-miRNA and target RNA (Gregory et al., 2005; Maniataki and Mourelatos, 2005). This process is heavily dependent on Ago-Dicer1 interaction (Bose and Bhattacharyya, 2016; Filipowicz et al., 2008). We have noted a marked loss of Ago2-Dicer1 interaction in lipid treated Huh7 cells, implicating impaired assembly of miRLC. Since, miRNA biogenesis is coupled to RISC activity, loss of Ago2-Dicer1 interaction causes reduced loading of miR-122 to Ago2 rendering reduced functional miR-122 specific miRNP formation. Concomitantly, Dicer1 processivity decreases as its active site remains associated with mature miR-122 and hence, it cannot bind a new pre-miR-122 molecule for processing. This is evident from increased association of NH-Dicer1 with miR-122 and miR-122*, together with reduced association of NH-Dicer1 with pre-miR-122 in lipid loaded Huh7 cells. Stability and processivity of Dicer1 is known to be governed by its interaction with other factors (Chendrimada et al., 2005; Haase et al., 2005). Thus, observed translocation of Dicer1 from ER to MVBs upon lipid treatment could be attributed to its reduced interaction with Ago2 which might lead to alteration in its stability, post-translational modifications or interaction with other factors that ultimately aid in MVB packaging of Dicer1.

Recent developments in the field of lipid biology, particularly studies on lipid droplets (LDs), have led us to the understanding of a wide range of functions of LDs. LDs also perform a wide range of non-canonical functions other than lipid storage (Welte, 2007) and are known to have contact sites with every other organelle. Owing to its dynamic nature, LDs are speculated to be potential vehicles for inter-organellar trafficking of various lipids and non-lipid factors inside the cell (Olzmann and Carvalho, 2019; Pennetta and Welte, 2018). Proteomic analysis and functional studies have identified a plethora of proteins associated with LDs which regulate various metabolic and signalling pathways. Most of the proteins targeted to LDs remain attached to it via amphipathic helices and hydrophobic hairpins (Thiam et al., 2013). There is a dynamic exchange of protein factors between the LD and its surroundings. Often LDs act as sequestration sites for protein function inactivation, for example the storage of histone proteins (Li et al., 2012) or MLX family of transcription factors (Mejhert et al., 2020). Interestingly, we observed microscopically a strong interaction of LDs with Ago2 in lipid challenged Huh7 cells. Our *ex vivo* and *in vitro* data indicate a role of LDs and their associated factors in regulation of Ago2-Dicer1 interaction. This is accredited to the sequestration of Ago2 by LDs and thereby, decreasing the active pool of Ago2 available for interaction with Dicer1 and subsequent miRNA loading. A recent study suggests that LDs possess phase behaviour and undergo liquid-crystalline phase transitions depending upon the ratio of triacylglycerol and cholesteryl ester. Such phase transitions regulate their interaction with other organelles and thus, have a functional significance (Mahamid et al., 2019). The possibility that LDs could phase out Ago2 from the cytoplasm may thus be a unique way of buffering miRNA activity in hepatic cells.

Liquid-liquid phase separation is the basis for formation of membrane less organelles. These phase separated condensates provide an alternative means of compartmentalization in cells. The condensates have been shown to be formed by phase separation of scaffold proteins (Banani et al., 2017). The inclusion/exclusion of a factor from a condensate depends upon the miscibility and energetically favourable interaction of the factor in the condensate, which in turn is governed by regulation of various modifications of the factor (Wheeler and Hyman, 2018). This mechanism thus provides the basis for a highly dynamic structure that undergoes rapid exchange of factors with the surrounding cytoplasm (or nucleoplasm) depending upon various cellular and environmental cues. P-bodies are membraneless phase separated entities that exhibit dynamic dissolution/condensation for localization at different cellular sites often in close proximity with endo-lysosomal structures (Liao et al., 2019). P-bodies comprise of several RNA degrading enzymes and are known to host repressed mRNAs which respond to cellular stress and mobilize reversibly from translating polysomes to P-bodies and *vice versa*. The miRNA effector protein Ago2 has also been reported to localize inside cytoplasmic P-bodies under certain circumstances (Filipowicz et al., 2008; Patranabis and Bhattacharyya, 2018).

In the nucleus of hepatocyte derived cell lines, LDs have been found to co-localize with the phase separated premyelocytic leukemia (PML) bodies and this interaction is necessary for the generation and stability of nuclear LDs (Ohsaki et al., 2016). We report a similar interaction of LDs with a cytoplasmic phase separated organelle, P-bodies in lipid loaded Huh7 cells. This interaction possibly facilitates exchange of specific factors between the P-bodies and LD surface. Interestingly, we observed a selective partitioning of Ago2 proteins from P-bodies to LDs (as calculated from relative co-localization coefficients of Ago2 with both the structures). The translocation of Ago2 to LDs might be promoted by certain post-translational modification which reduces the miscibility of Ago2 in P-body condensates. We also noted an uncoupling of miR-122 from Ago2 in presence of LDs. This can be attributed to certain modifications such as phosphorylation (Patranabis and Bhattacharyya, 2016; Patranabis and Bhattacharyya, 2018) of Ago2 happening at the P-body-LD interface. Uncoupling of miR-122 from Ago2 might also be due to the presence of some unidentified RNA binding proteins on the LD surface that have a higher affinity for miR-122 and thus, shifts the equilibrium from Ago2-miR-122 interaction to the miRNA free Ago2 population-a process which may also be accelerated by miRNA-binder HuR that reversibly binds miRNA (Mukherjee et al., 2016) to facilitate this phase separation. We detected elevated cellular levels of HuR in lipid challenged Huh7 cells. Additionally, HuR had a role in modulating cellular level of Dicer1 as depletion of HuR was found to inhibit the reduction in cellular Dicer1 levels in lipid treated Huh7 cells. HuR knockdown also corroborated with rescue in cellular miR-122 levels in lipid loaded Huh7 cells (data not shown). Earlier reports suggested the involvement of HuR in binding miR-122 and facilitating MVB packaging and export of miR-122 inside EVs from starved Huh7 cells (Mukherjee et al., 2016). Interestingly, we detect enrichment of NH-Dicer1 in EVs isolated from HA-HuR expressing Huh7 cells as compared to control Huh7 cells. Studies showed that HuR not only interacts with miR-122, but also interacts with miR-122*, thus, implicating its probable role in interaction with Dicer1 and influencing miRNA biogenesis as well (Mukherjee et al., 2016). Surprisingly, we find that ectopic expression of HA-HuR causes a loss of Ago2-Dicer1 interaction in Huh7 cell. It would be interesting to look at Dicer1-HuR interaction in lipid treated Huh7 cells and whether such an interaction directly leads to loss of Ago2-Dicer1 interaction in Huh7 cells.

## Materials and Methods

### Cell Culture and transfections

Huh7.and HEK293, were cultured in Dulbecco’s Modified Eagle’s Medium (DMEM) supplemented with 2mM L-glutamine and 10% heat inactivated fetal calf serum and 1% Penicillin-Streptomycin.

All plasmid transfections to Huh7 and HEK293 cells were done with Lipofectamine 2000 (Invitrogen), according to the manufacturer’s protocol. 1µg of plasmid was transfected per well of a six-well plate. siRNAs were transfected using RNAiMAX (Invitrogen), according to the manufacturer’s protocol. 30 picomoles of respective siRNAs were transfected per well of a twelve-well plate.

### Cholesterol and Palmitic Acid treatment

MβCD conjugated cholesterol conjugate obtained from GIBCO (#12531-018) was added from a 250X stock to Huh7 cells in culture at a final concentration of 1X, 2X or 5X for a period of 2, 4 or 8 hours. Cholesterol treatments were done to Huh7 cells in fresh growth media at 70-80% confluency. Unless otherwise mentioned, cholesterol treatment was done at a final concentration of 5x for 4 hours.

Palmitic acid (Sigma) was conjugated to fatty acid free BSA (Sigma) in KRBH buffer (NaCl 135mM, KCl 3.6mM, NaHCO_3_ 5mM, NaH_2_PO_4_ 0.5mM, HEPES 10mM, MgCl_2_ 0.5mM, CaCl_2_ 1.5mM, D-glucose 3.5 mM, BSA 21%) at 37°C for 6 hours in a shaking condition. Final concentration of this working stock is 10mM with a BSA:PA (Palmitic Acid) ratio of 1:3. The BSA conjugated Palmitic acid was then filter sterilized and added to Huh7 cells at a final concentration of 300, 500 or 1000µM for 16 hours. For control (0 µM), equal amount of fatty acid free BSA in KRBH buffer was added. All treatments were done to Huh7 cells in fresh growth media at 50-60% confluency.

### Statin Treatment

For inhibition of HMG-CoA reductase, Huh7 cells were cultured in serum free Dulbecco’s modified Eagle’s medium (DMEM; Gibco-BRL) for 24 h. Atarvostatin stock concentration of 1mg/ml (1.78 mM) was dissolved in UltraPure distilled water and added to the cells at concentrations of 0, 1, 5 and 10 µM. Cells were subsequently incubated over a time period of 48 hours.

### Western Blotting

Cell lysates or immunoprecipitated protein extracts were subject to SDS-polyacrylamide gel electrophoresis followed by transfer of proteins to a polyvinylidene difluoride (PVDF) membrane (Milipore). The membranes were blocked for 1 hour in blocking buffer containing Tris-buffered saline (TBS) containing 0.1% Tween 20 and 3% BSA at room temperature.

The blots were then hybridized with primary antibodies in blocking buffer for a minimum time of 16 hours at 4°C. This was followed by washing of the blots thrice for 5 min each in TBS containing 0.1% Tween 20. The blots were then probed with secondary antibodies (1:8000 dilution) for 1 hour at room temperature. Excess antibodies and non-specific bindings were washed off with TBS-Tween 20 thrice for 5 min each. Development and imaging of all the western blots signals was done in UVP Bioimager 600 system equipped with VisionWorks Life Science software (UVP) V6.80. Quantification of band intensities was done using ImageJ software.

### RNA isolation and quantification of miRNA and mRNA levels

RNA isolation was done using TRIzol reagent (Life Technologies) according to the manufacturer’s protocol. Real-time quantitative RT-PCR of mRNA was done with 100-200 ng of RNA by a two-step reaction format using Eurogentec Reverse Transcriptase Core Kit and MESA GREEN qPCR Master Mix Plus for SYBR Assay-Low ROX. 18S ribosomal RNA was used as an endogenous control. For expression of lipid metabolic genes, β-Actin mRNA was used as an endogenous control.

Real-time quantitative RT-PCR of miRNA, namely miR-122, miR-16, miR-21 and other miRNAs was done with 25 ng of total RNA using Applied Biosystems TaqMan chemistry based assays, following the manufacturer’s instructions. U6 snRNA was used as the endogenous control. For detection of miRNAs in extracellular vesicle fractions, 100ng of RNA was used for reverse transcription in each cases.

All PCR reactions were done in either 7500 Applied Biosystem Real Time System or BIO-RAD CFX96 Real-Time system. The RT reaction conditions were 16°C for 30 min, 42° C for 30 min, 85°C for 5 min, followed by product held at 4°C. The PCR conditions were 95°C for 5 min, followed by 40 cycles of 95°C for 15 seconds, 60°C for 1 min.

### Extracellular Vesicle (EV) isolation

For all EV related experiments cells were grown in media supplemented with EV free FCS. EV depleted FCS was either commercially obtained (System Biosciences Catalog no. EXO-FBS-250A-1) or prepared by ultracentrifugation of the FCS used at 120,000 x g for 5 hours. For EV isolation, the supernatant conditioned media (CM) from two 60 cm^2^ plates, having 6×10^6^ donor cells (Huh7) each were taken. CM were pre-cleared by centrifugation at 300 x g for 10min, 2,000 x g for 15 min, 10,000 x g for 30 min at 4° C and subsequently, filtered through a 0.22 um filter unit. The resultant filtrate was then loaded on top of a sucrose cushion (1 M sucrose and 10 mM Tris–HCl pH 7.5) and ultracentrifugation was done at 120,000 x g for 90 min at 4° C. The media above the sucrose cushion was discarded till the interface leaving behind a narrow layer of medium where the EVs get enriched. The EVs where then washed with PBS by ultracentrifugation at 100,000g for 90 min at 4° C. The pellet was resuspended in 200 µl 1X Passive Lysis Buffer (Promega) and divided into two parts for protein and RNA analysis, respectively.

### Subcellular fractionation on OptiPrep^TM^ gradients

Cell fractionation was done on a OptiPrep^TM^ gradient as previously described (Chakrabarty and Bhattacharyya, 2017). Briefly, a 3-30% or 3-15.5% OptiPrep^TM^ (Sigma) continuous gradient was prepared in a buffer containing 78 mM KCl, 4 mM MgCl_2_, 10 mM EGTA, 50 mM HEPES pH 7.0. Cells were harvested and washed in ice-cold PBS followed by homogenization using a Dounce homogenizer in a buffer containing 0.25 M sucrose, 78 mM KCl, 4 mM MgCl2, 8.4 mM CaCl_2_, 10 mM EGTA, 50 mM HEPES pH 7.0 alongwith 100 ug/ml Cycloheximide, 0.5 mM DTT, 40 U/ml RNase Inhibitor (Applied Biosystems), 1X Protease inhibitor (Roche). The lysate was cleared by centrifugation at 1,000g for 5 min, twice. The clarified lysate was loaded on top of the OptiPrep^TM^ gradient and ultracentrifuged at 36,000 rpm for 5h using Beckman Coulter SW60Ti rotor at 4° C. After ultracentrifugation, ten fractions were collected by aspiration from top of the tubes. The fractions were then processed for RNA or protein analysis.

### Lipid Droplet isolation

Lipid droplets (LDs) were isolated from Huh7 cells by floatation gradient ultracentrifugation according to the published protocol (Ding et al., 2013) with minor modifications. Briefly, fatty acid treated or untreated Huh7 cells were homogenized using a Dounce homogenizer in Buffer A [containing 20 mM Tricine, 250 mM sucrose, 1X protease inhibitor, 40 U/ml RNase inhibitor (Applied Biosystems), pH 7.8]. The homogenate was clarified by centrifugation at 1,000xg for 5 min, twice. The clear post-nuclear supernatant (PNS) was loaded into a SW60Ti compatible ultracentrifugation tube. A layer of Buffer B [containing 20 mM HEPES, 100 mM KCl, 1X protease inhibitor, 40 U/ml RNase inhibitor, pH 7.4] was carefully loaded on top of the PNS and the resultant gradient was ultracentrifuged at 31,200 rpm for 1h and 15 min in a Beckman Coulter SW60Ti rotor at 4° C. Post ultracentrifugation, LDs were collected from the top band of the gradient, cytosolic fraction was collected from the middle part of the gradient, and total membrane fraction was the pellet. The obtained LDs were washed thrice in Buffer B by centrifugation at 20,000xg for 5 min at 4° C.

### Microsome isolation

Microsomes were isolated from Huh7 cells as previously described (Barman and Bhattacharyya, 2015). Briefly, Huh7 cells were washed with PBS and resuspended in 1X hypotonic buffer (containing 10 mM HEPES, pH 7.8, 1 mM EGTA, 25 mM KCl) proportionate to three times packed cell volume for 20 min on ice. The cells were then resuspended in 1X isotonic buffer (10 mM HEPES, pH 7.8, 1 mM EGTA, 25 mM KCl, 250mM sucrose) proportionate to two times packed cell volume, followed by homogenization. The homogenate was clarified by centrifugation at 1000g for 5 min. The supernatant was centrifuged at 12,000xg for 10 mins in order to remove the mitochondrial fraction. The post-mitochondrial supernatant was incubated with 8M CaCl_2_ for 30 min at 4° C in a rotator, followed by centrifugation at 8,000xg for 10 min to get the microsomal pellet.

### Immunoprecipitation

Immunoprecipitation of specific proteins was carried out according to the published method (Bhattacharyya et al., 2006). Briefly, cells were lysed in a lysis buffer [containing 20mM Tris-HCl pH 7.4, 150mM KCl, 5mM MgCl_2_, 1 mM dithiothreitol (DTT), 1X EDTA-free protease inhibitor (Roche), 40 U/ml RNase Inhibitor (Applied Biosystems), 0.5% Triton X-100, 0.5% sodium deoxycholate] for 30 min at 4° C. The lysate was cleared by centrifugation at 3,000xg for 10 min. Protein G Agarose beads (Invitrogen) were blocked with 5% BSA in lysis buffer for 1h. The bead were then allowed to bind to the required antibody for 3-4 h at 4°C in lysis buffer, followed by addition of the lysate. A final dilution of 1:100 (anibody:lysate) was used for immunoprecipitation. All immunoprecipitations were done for 16 h at 4°C followed by bead washing thrice in IP Buffer [containing 20mM Tris-HCl pH 7.4, 150mM KCl, 5mM MgCl_2_, 1 mM dithiothreitol (DTT), 1X EDTA-free protease inhibitor (Roche), 40U/ml RNase Inhibitor (Applied Biosystems)]. The beads were then divided into two parts; one part for RNA isolation and the other part for western blotting.

### *In vitro* assays with LD

*In vitro* Dicer1-Ago2 interaction assay in presence or absence of LDs was done as follows. Briefly, NH-Dicer1 expressing HEK293 cells were homogenized in a Dounce homogenizer in Buffer A. The homogenate was clarified by centrifugation at 1,000xg for 5 min, twice. The clarified homogenate was incubated with LDs isolated from Palmitate stimulated Huh7 cells at 37°C for 1h. For control, equal amount of Buffer B was added to the clarified homogenate. Post incubation, immunoprecipitation of NH-Dicer1 was done using α-HA antibody at 4° C for 16 h, according to the protocol already mentioned above. The resulting sample was then analysed by Western blotting.

For RNase protection assay of LD associated miR-122, LDs from Palmitate stimulated Huh7 cells were initially isolated in Buffer B1 [containing 20 mM HEPES, 100 mM KCl and 2 mM KCl, 1X protease inhibitor, pH 7.4]. RNase digestion of the LD associated miR-122 was carried out by incubation with RNase A (Fermentas). RNase digestion was done in 100 µl of reaction volume containing isolated LDs, RNase A (10 µg or 20 µg/ml) in Buffer B1 at 30° C for 1 h. RNA extraction was done with TRIzol LS, followed by RNA precipitation at -20° C for 4 h. Isolated RNA was analysed for quantification of miR-122 by qRT-PCR.

For in-vitro uncoupling of miR-122 from Ago2 in presence of LDs, HEK293 cells expressing FH-Ago2 and miR-122 were homogenized with a Dounce homogenizer in Buffer A. The homogenate was clarified by centrifugation at 1,000xg for 5 min. The clarified homogenate was incubated with LDs (isolated from Palmitate stimulated Huh7 cells) for increasing time points, followed by immunoprecipitation of FH-Ago2 (with anti-HA antibody) for 16h at 4°C. The immunoprecipitated extract was then analysed for the amount of FH-Ago2 pulled down (by Western blot) and amount of miR-122 associated with it (by qRT-PCR)

### Immunofluorescence Microscopy and post-imaging analysis

Cells grown on gelatin coated 18 mm round coverslips were transfected and treated as indicated. Cells were then fixed with 4% para-formaldehyde in PBS for 30 min at room temperature in dark. For immunostaining, blocking and permeabilization was done in 1% BSA in PBS containing 0.1% Triton X-100 and 10% goat serum (Gibco) for 30 min at room temperature followed by probing with appropriate primary antibodies for 16h at 4°C. The coverslips were washed thrice in PBS followed by staining for 1h at room temperature with Alexa Fluor® anti-rabbit, anti-mouse or anti-rat secondary IgG conjugated to appropriate fluorochrome (Moleclar probes). For lipid droplet staining, Bodipy 493/503 (Invitrogen) was used. Coverslips were mounted on cleaned glass slides using VECTASHIELD mounting medium (with DAPI). Confocal fixed-cell imaging was done with Zeiss LSM 800 confocal system. All post imaging analysis was done using Imaris 7 (BitPlane) software.

### Fluorescence Correlation Spectroscopy (FCS)

Single point FCS measurements on live cells were performed using an ISS Alba FFS/FLIM Confocal Microscope (Champaign, IL, USA) using a pulsed 488 nm diode laser excitation. We used a C Apochromat 63 X water immersion objective and cells cultured in Lab-Tek II chambered coverglasses (Nunc, Roskilde, Denmark) were imaged by CLSM methods. The fluorescence emissions at 532 nm were detected using a pair of SPAD (Single Photon Avalanche Detector) detectors and single color correlation functions were calculated to minimize the effects of after-pulsing. Specific region of interests (ROIs) were selected in the cell cytosol in which the FCS experiments were carried out. Acquisitions for 60 seconds each were carried out at the individual ROIs, and a collection of three independent sets of experiments was used for each correlation function.

For the analyses of the correlation functions, we initially used a single-component diffusion model as described by the following equation 1:

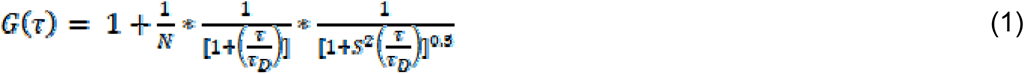

where, τ_D_ is the translational diffusion time of the fluorescently-labelled diffusing protein. N is the average number of protein molecules in observation volume. S defines the ratio of beam radius r_0_ to beam height z_0_ of the instrument. The parameter τ_D_ is related to the diffusion coefficient (D) as per the following equation 2:

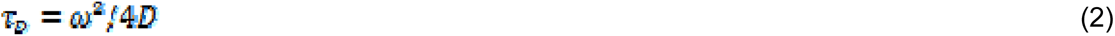

where ω is the size of the observation volume, the value of which was calculated using Rhodamine 6G as reported earlier (Chattopadhyay et al., 2005).

We found that the correlation functions obtained in our experiments could not be fit using equation 1. Considering the cell cytosol to be crowded, our data were fit to an anomalous diffusion model given by following equation 3 as reported earlier (Banks and Fradin, 2005):

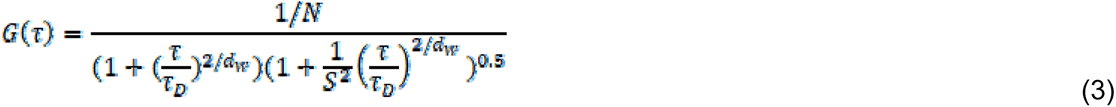

All data analyses were carried out using Origin Pro (OriginLab Corp., MA, USA).

### Statistical Analysis

All graphs were plotted and analysed using GraphPad Prism 5.00 (GraphPad, San Diego, CA, USA). For statistical analysis, non-parametric paired student t-test was performed. Error bars indicate mean with standard deviation.

### Animal experiments

All animal experiments were approved by the Institutional Animal Ethics Committee (approved by CPCSEA, Ministry of Environment & Forest, and Government of India). 8-10 weeks old male (20-24g) C57BL/6 mice were housed under controlled conditions (temperature 23 ± 2c, 12 hour/12-hour light/dark cycle) in individually ventilated cages. Mice were randomly divided into two groups and fed either standard chow diet or methionine and choline deficient diet MCD (MP Biomedicals; #0296043910) up to four weeks.

C57BL/6 male mice of 8-10 weeks age were divided in two groups for normal chow and high fat diet (HFD) containing 45% fat and 5.81 kcal/gm diet energy content (MP Biomedicals; # 960192). Animals were fed with HFD for 4 weeks.

For isolation of RNA from tissues, TRIzol® (Invitrogen) reagent was used. For analysis of EV-associated RNA, serum fraction of blood was used. Relative levels of miRNA and mRNA in serum and tissues were quantified by qRT–PCR. For analysis of proteins, liver slices were homogenized in 1× RIPA buffer (25 mM Tris–HCl pH 7.6, 150 mM NaCl, 1% NP-40, 1% sodium deoxycholate, 0.1% SDS) to generate lysates. Lysates were prepared by sonication followed by clearing with centrifugation at 20,000 × *g*, at 4°C for 1 h, and relative levels of endogenous Dicer1 was detected by Western blot analysis using anti-Dicer1 antibody. 50 μg of protein was loaded per well of SDS PAGE analysis to detect Dicer11. β-Actin was used as the loading control.

For histological analysis, tissues were fixed in 10% formaldehyde in PBS, embedded in paraffin, sectioned at 10 μm, and stained with hematoxylin and eosin (H&E) following standard staining protocol.

For detection of serum miRNA levels, serum fraction of blood was used. Blood samples were collected by cardiac puncture and allowed to clot. Serum was separated by centrifugation and frozen at -80°C.

### Primary Hepatocyte Isolation

Animals were obtained from the animal house of the institute and all experiments were performed according to the guidelines set by Institutional Animal Ethics committee following the Govt. of India regulations. Mouse primary hepatocytes were isolated using the hepatocyte product line from Gibco Invitrogen Corporation. Adult BALB/c mice (4-6 weeks) were anaesthetized and the portal vein was cannulated using a 25G butterfly cannula and an incision was made in the inferior vena cava. The liver was perfused with 350 mL of warm (37°C) Liver Perfusion Medium (Cat. No. 17701) at a rate of 35 mL/minute with the perfusate exiting through the severed vena cava. This was followed by a Collagenase-Dispase digestion with Liver Digest Medium (Cat no. 17703) at a rate of 35 mL/minute. The liver was then aseptically transferred to the tissue culture hood on ice in Hepatocyte Wash Medium (cat no. 17704). Using blunt forceps the digested liver was torn open to release the hepatocytes. Cell clumps were dissociated by gently pipetting the solution up and down using a 25ml pipette. The solution was then filtered through 100 μM nylon caps atop 50 ml conical tubes. The cell suspension was then centrifuged at 50 x g for 3 min. The pellet was gently resuspended in 10 ml of Wash Medium using 25 ml pipette and the centrifugation repeated.

Cells were finally resuspended in Hepatocyte Wash Medium with 10% FCS and plated at 1×10^7^ cells /ml. Cells were plated in tissue culture treated collagen (Gibco Cat. No. A10483-01) coated plates at 12.5 µg/cm^2^. Unattached cells are poured off 4h after plating and medium was replaced with Hepatozyme-SFM (Cat no. 17705) with glutamine and 1% Pen/Strep. Cholesterol and BSA-Palmitate was added the next day in Hepatozyme-SFM.

### TUNEL Assay

TUNEL assay was performed using Invitrogen Click-iT^™^ TUNEL Alexa Fluor 594 Imaging Assay kit for HA-DICER1 expressing Huh7 cells, as per manufacturer’s protocol.

### Differential expression analysis and pathway mapping

In order to determine alterations in hepatic gene expression profile under lipotoxic stress and study the possible physiological relevance of expression changes in miR-122 target genes, differentially expressed genes and their pathway involvement has been studied. Differential expression analysis was performed considering high fat diet fed mouse versus chow diet fed mouse for 4 weeks, 12 weeks and mice having high fat along with cholesterol diet as opposed to only high fat diet for 16 weeks. Samples from GSE53381 [high fat diet treatment for 4 weeks (HFD4)] and GSE93819 [high fat diet treatment for 12 weeks (HFD12)] (Kobori et al., 2017)) datasets were considered to identify genes that are involved in regulating short term or long term high fat diet exposure associated stress. Additionally, samples from GSE58271 (Lorbek et al., 2015)) were considered to identify genes involved in responding to high cholesterol (CH) exposure associated stress. For this purpose, differentially expressed genes were determined with the help of limma (Ritchie et al., 2015) R package in Gene Expression Omnibus (GEO) series datasets ((Barrett et al., 2013)). Genes having fold change >= 1.5 and p-value <= 0.05 were considered as differentially expressed and are likely to be involved in regulating lipotoxic species associated stress. A probable set of miR-122 target mRNA was determined based on a previous study (Wen and Friedman, 2012) wherein livers of miR122a^-/-^ mice had been compared with wild-type mice. mRNA that had fold change in expression higher than 1.5 or lower than -1.5 with significant p-values have been considered as miR-122 target mRNAs. Venn diagrams were utilised to determine miR-122 target genes that showed altered levels under two or more of these conditions. Additionally, pathway mapping of miR-122 target genes differentially expressed in two or more of the high fat diet treatment conditions was performed. The genes were mapped onto KEGG database (Cokelaer et al., 2013) pathways with the help of a python script. Cellular pathways belonging to the categories such as ‘carbohydrate metabolism’, ‘lipid metabolism’, ‘cellular process’ and ‘signal transduction and signalling molecules and interaction’ were considered for the analysis. A plot was prepared to determine whether the high fat diet exposure associated miR-122 target genes participated in multiple signalling or metabolic pathways and as such whether alterations in miR-122 target genes may bring about time dependent physiological changes. miR-122 target genes that exhibited high fat diet exposure associated alterations were represented in rows with their corresponding pathway involvement(s) being shown in columns and each cell depicted the conditions under which they exhibit alterations.

Information on Plasmids, Oligos, PCR primers, Antibodies are provided in Supplementary Table 1-6.

## Acknowledgements

We thank Witold Filipowicz and Gunter Meister for different constructs used in this study. We thank the Funding body, Dept. of Science and Technology (DST), Govt. of India and Council for Scientific and Industrial Research (CSIR), and University Grant Commision (UGC) for the fellowship to DB, SB, IM, RC,KM and MA. SB also acknowledges the support from Department of Biotechnology, Govt of India for her Fellowship support. SNB was supported by SwarnaJayanti Fellowship and a High Risk High Reward Grant from DST.

## Author Contributions

S.N.B. conceived the idea, designed the experiments, analyzed the data and wrote the manuscript. K.M, S.B. S.C. K.C. and P.C have contributed in design and planning the experiments. D.B., S.B., I.M., R.C., M.A. and K.M. performed the experiments. S.B. and D.B also wrote the manuscript with S.N.B. and analyzed the data.

## Supplementary Figure Legends

**Supplementary Figure S1.**
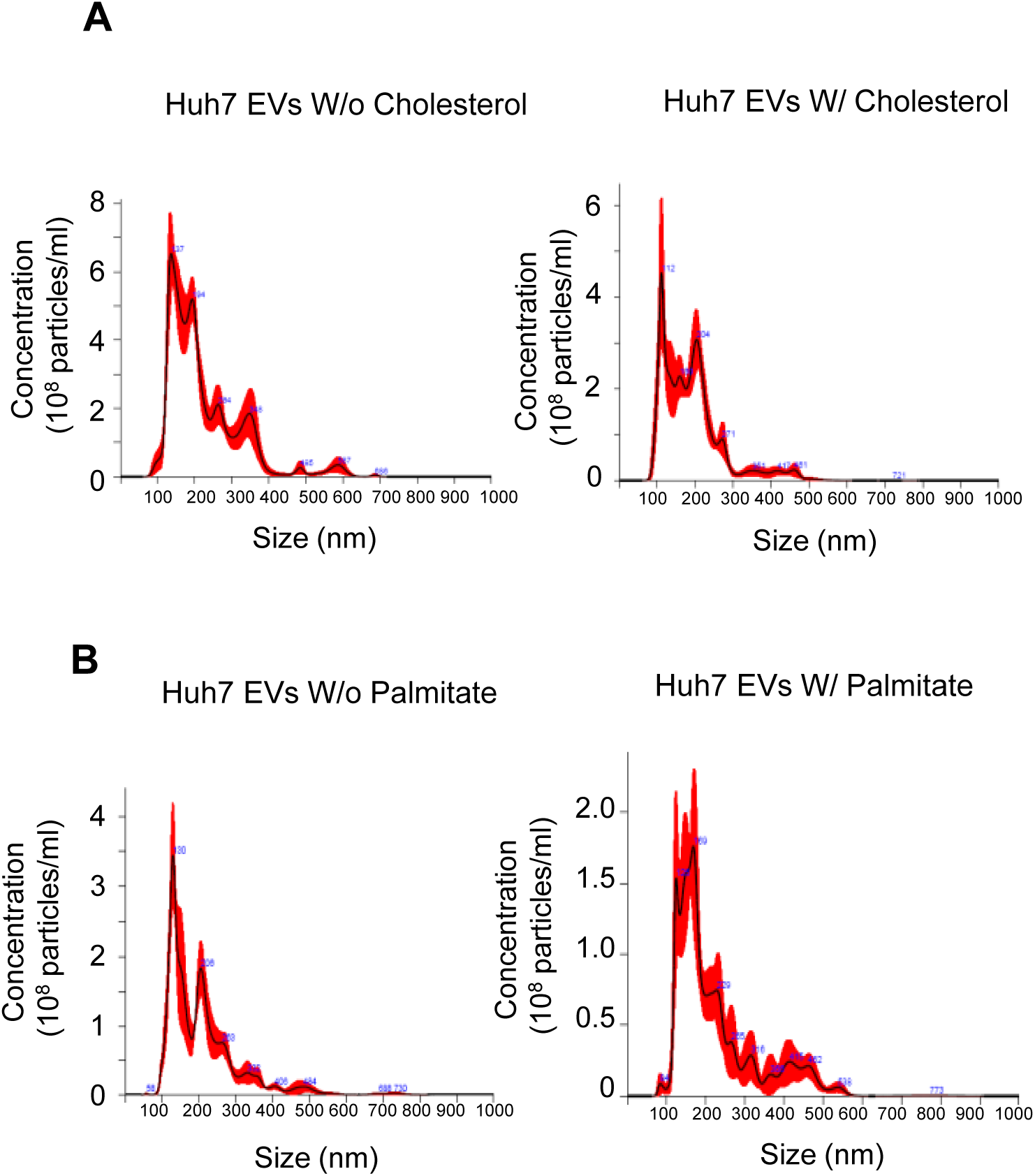
Nanoparticle tracking analysis of the EVs isolated from control and lipid exposed cells. **A-B** Representative graphs from Nanoparticle Tracking Analysis (NTA) depicting the size vs. concentration profile of isolated EVs from control, cholesterol treated (A) and BSA-Palmitate treated (B) Huh7 cells. NTA graph of EVs isolated from culture supernatants of control Huh7 cells (panel A, left), cholesterol-lipid treated Huh7 cells (panel A, right), fatty acid free BSA treated Huh7 cells (panel B, left) and BSA-Palmitate treated Huh7 cells (panel B, right) are shown.

**Supplementary Figure S2.**
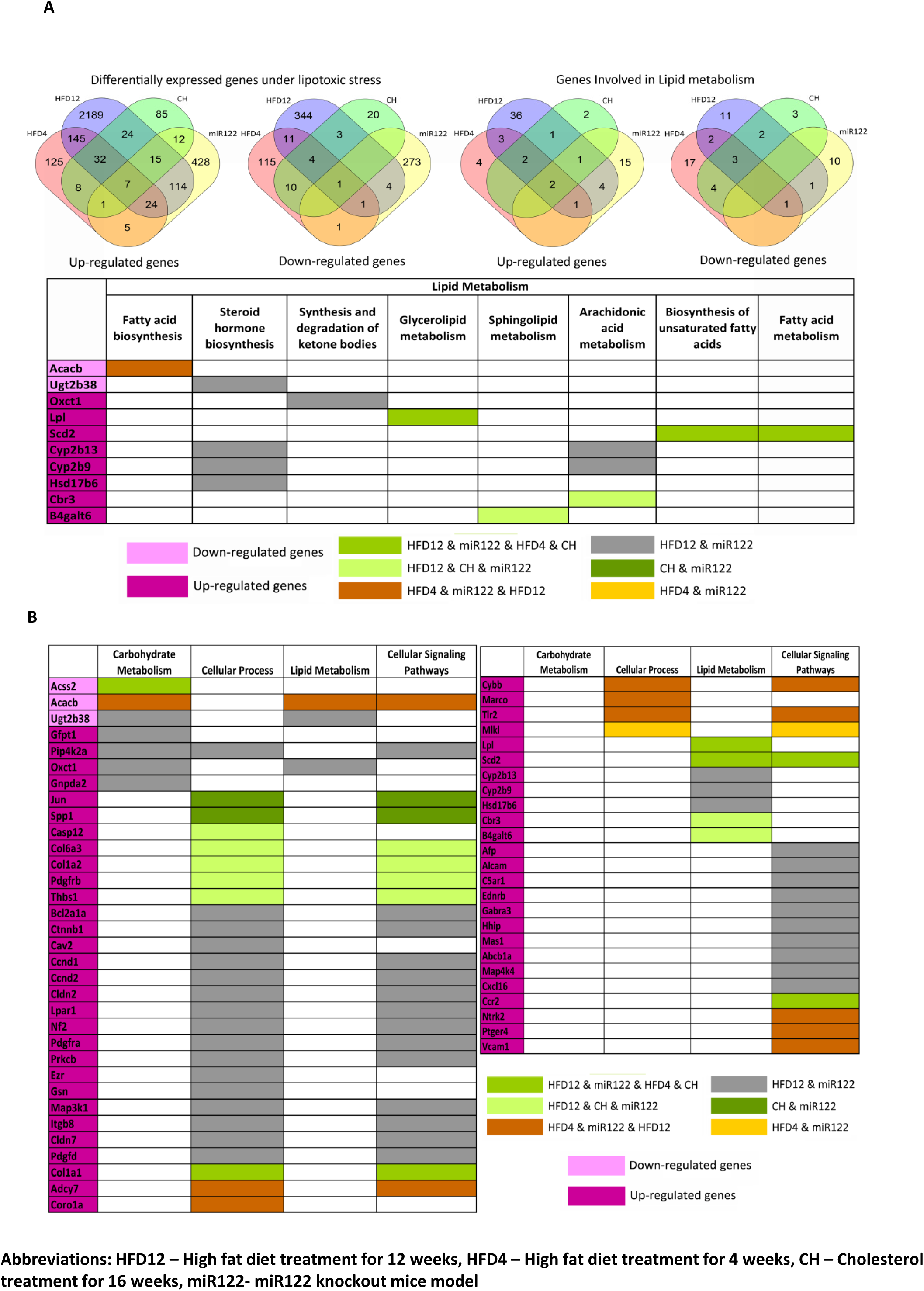
Differentially regulated genes and possible de-regulated cellular pathways on high fat diet treatment or miR122 ^(-/-)^ in mice models. **A** Hepatocyte cells in high fat diet treated mouse models exhibit alterations in expression levels of miR-122 target genes. A comparison of genes de-regulated in various High Fat Diet case studies (HFD12, HFD4, CH) has been shown here. Genes are mainly up-regulated upon high fat diet treatment and the overlap among these genes and miR122 targets is shown in (first from left, top panel). Down-regulated genes that occur upon high fat/cholesterol treatment and upon miR-122 knock-out in mice models is also shown here (second from left, top panel). Lipid metabolism associated miR-122 target mRNA that were found to be up-regulated upon exposure to high fat conditions for different time periods (third from left, top panel) and number of lipid metabolism associated miR-122 target mRNA that become down-regulated under high fat diet treatment conditions in mice models is shown (fourth from left, top panel). Differentially expressed miR-122 target genes that are involved in regulating lipid metabolism under conditions of lipotoxic stress induced due to high fat diet treatment (bottom panel). **B** Differentially expressed miR-122 target genes under lipotoxic stress that are involved in multiple pathways are shown here. Down-regulated genes are associated with carbohydrate and lipid metabolism whereas up-regulated genes are mainly associated with cellular signalling or cellular process associated pathways. Abbreviations, HFD12 – High fat diet treatment for 12 weeks, HFD4 – High fat diet treatment for 4 weeks, CH – Cholesterol treatment for 16 weeks, miR122- miR122 knockout mice model.

**Supplementary Figure S3.**
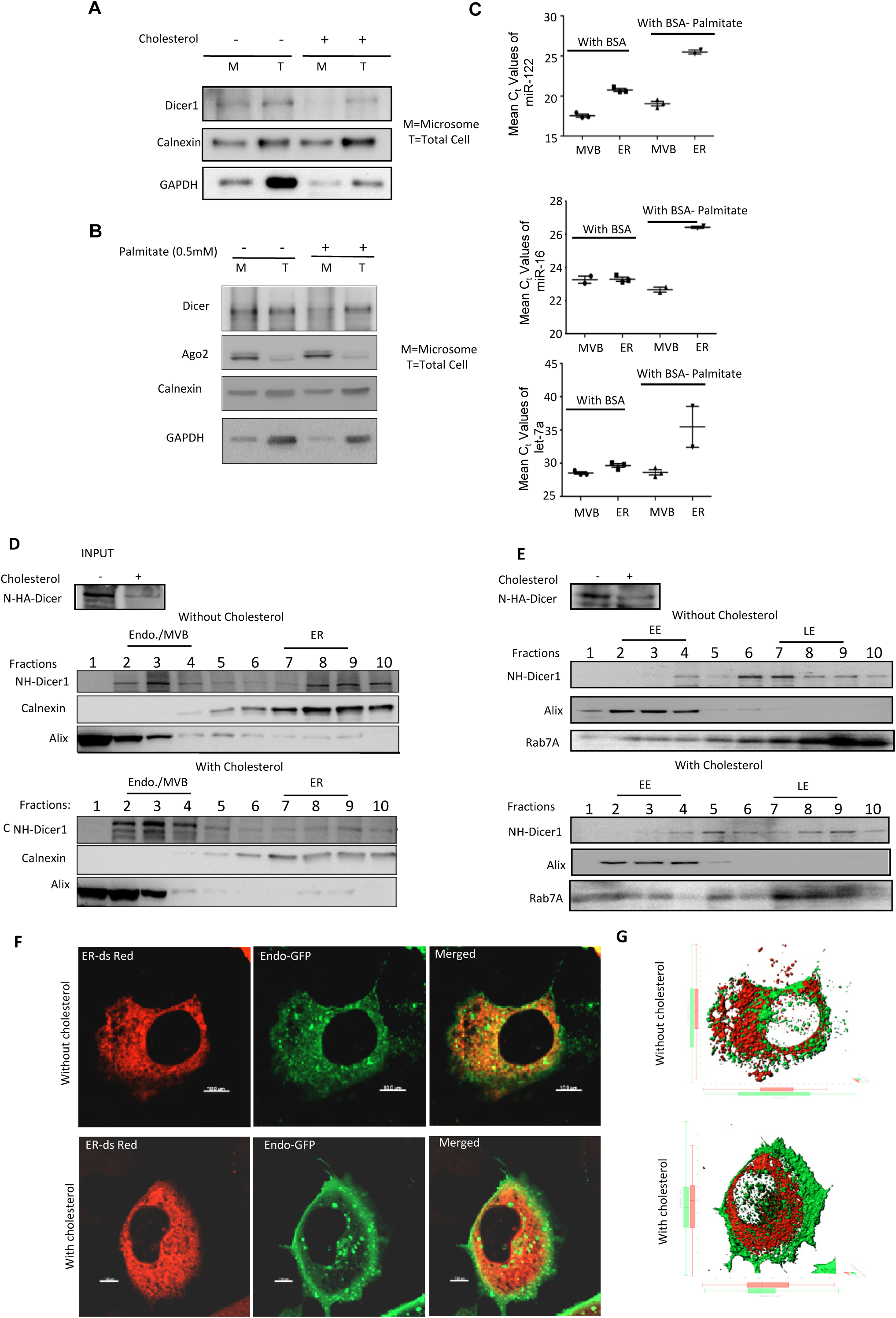
Cellular distribution of Dicer1in cholesterol treated cells. **A-B** Relative amount of Dicer1 and Ago2 in total and microsomal fractions of control, 5x cholesterol-lipid treated and BSA-Palmitate treated Huh7 cells. Microsomal fractions were isolated from cholesterol-lipid concentrate treated and untreated Huh7 cells. 10 µg of protein from total and microsomal fractions were analysed by western blotting to detect levels of Dicer1. Calnexin was used as a marker for the ER and GAPDH was detected to determine the level of cytosolic contamination (A). Also, microsomes were isolated from control (BSA treated) and 0.5mM BSA-Palmitate (16h) treated Huh7 cells. 10µg of protein from total and microsomal fractions were western blotted to detect levels of Dicer1 and Ago2 (B). Calnexin was used as marker for ER. GAPDH was blotted to check for cytosolic contamination. **C** Subcellular distribution of different miRNAs (miR-122, miR-16 and let-7a) in control and 0.5mM BSA-Palmitate treated Huh7 cells. Huh7 cells, transiently expressing NH-Dicer1 were either BSA treated or treated with 0.5mM BSA-Palmitate for 16h followed by lysis in isotonic conditions. Lysates were analyzed on a 3-30% iodixanol gradient for separation of organelles.. RNA was isolated from pooled fractions 2,3,4 (representing MVB) and pooled fractions 7,8,9 (representing ER) and qRT-PCR detection of miRNAs were done from volume equivalent samples. Ct values of the respective samples are plotted. **D** Subcellular distribution of NH-Dicer1 in Huh7 cells. Huh7 cells transiently expressing NH-Dicer1 were treated with or without 5x cholesterol-lipid concentrate for 4 hours. Cells were lysed in isotonic buffer and post-nuclear supernatants were subject to ultracentrifugation on a 3-30% iodixanol gradient for separation of subcellular organelles. Calnexin and Alix were used as markers of endoplasmic reticulum (ER) and endosomes/multivesicular bodies (MVBs), respectively. Positions of the respective organelles are marked on top of the blots. Western blot of NH-Dicer1 levels in control and 5X cholesterol-lipid concentrate treated Huh7 cells are shown as input. **E** Endosomal distribution of NH-Dicer1 in Huh7 cells studied by resolving of early and late endosomes/MVBs on 3-15% iodixanol gradient. Huh7 cells transiently expressing NH-Dicer1 were treated with or without 5x cholesterol-lipid concentrate for 4 hours. Cells were lysed in isotonic conditions and post-nuclear supernatants were ultracentrifuged on a 3-15% iodixanol gradient for separation of early and late endosomes/MVBs. The distribution of Alix and Rab7A shows the positions of early endosome and late endosome, respectively. Endosomal distribution of NH-Dicer1 was monitored by western blot with anti-HA antibody. Positions of the respective organelles are marked on top of the blots. Input levels of NH-Dicer1 are also shown. **F-G** Representative images depicting the distribution of ER and endosomes in control Huh7 and 5X cholesterol-lipid treated (4 hours) Huh7 cells (F). Huh7 were co-transfected with plasmids encoding an ER localizing variant of ds Red (ER-ds Red) and endosome localizing variant of GFP (Endo-GFP). ER-ds Red and Endo-GFP signals were detected by direct immunofluorescence (F). Fields have been detected at 60x magnification. Scale bar represents 10 µm. Panel (G) represents merged 3D surface reconstructions of ER and endosome, depicting their relative spatial distribution.

**Supplementary Figure S4.**
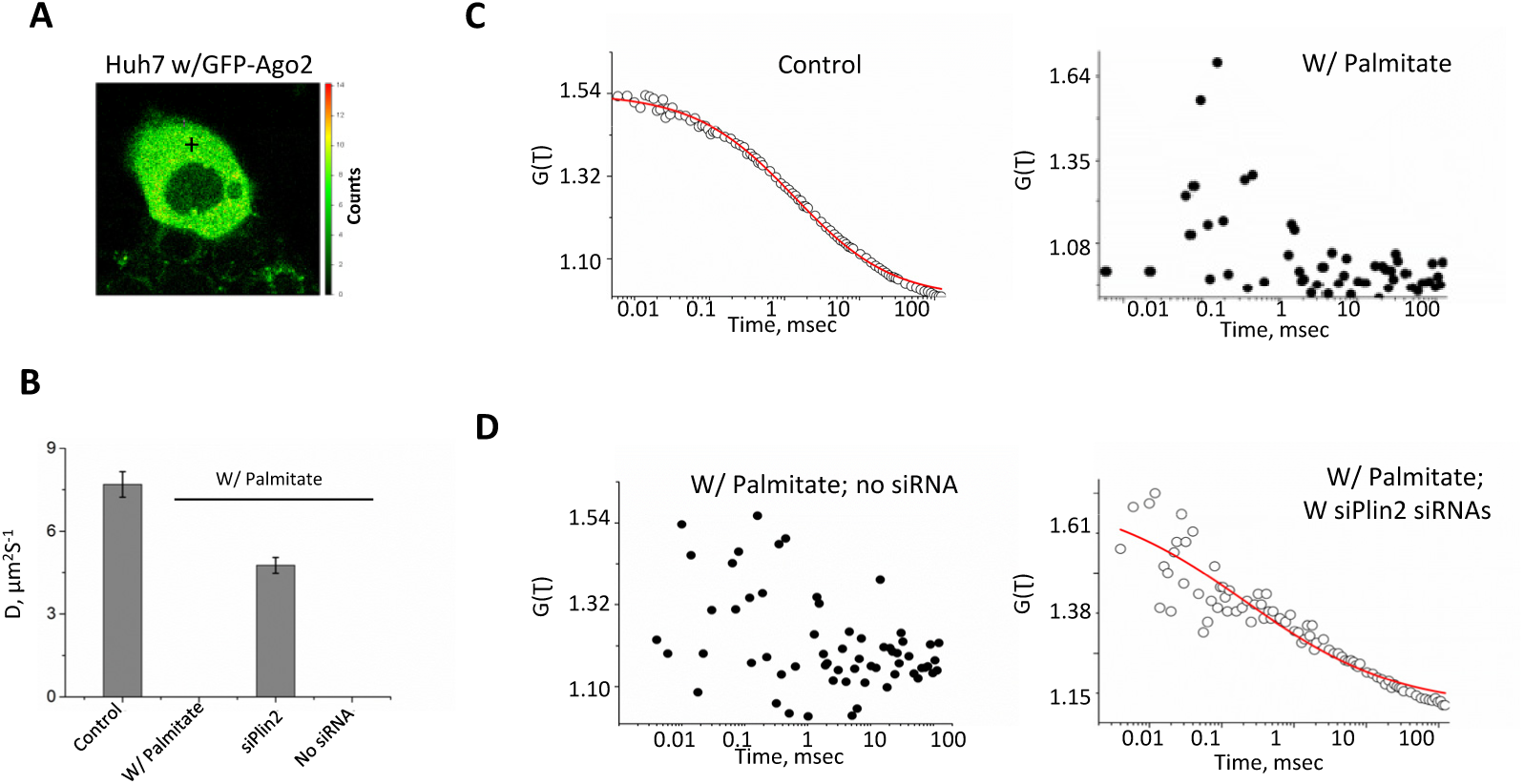
Reduced Ago2 dynamics in lipid exposed hepatic cells. **A-D** Fluorescence correlation spectroscopic analysis of GFP-Ago2 signals in control and BSA-Palmitate treated Huh7 cells, transiently expressing GFP-Ago2. Panel A shows a representative image with signal intensities of GFP-Ago2 molecules in Huh7 cells. Black cross mark indicates a representative position where single point FCS of GFP-Ago2 in the cytosol was performed. Autocorrelation curves of single point FCS measurements showing the diffusion pattern of GFP-Ago2 molecules in control (C, left panel), BSA-Palmitate treated (C, right panel), siControl and siPerilipin2 transfected, BSA-Palmitate treated Huh7 cells (D) have been plotted. The autocorrelation functions have been fitted to an anomalous diffusion model, considering that the cytosol is crowded. Variations of the diffusion coefficient of GFP-Ago2 in different experimental conditions are shown in panel B.

**Supplementary Table 1.** List of primary antibodies used

| Antigen name | Raised in | Source | Dilution for Western blot | Catalog no. |
| --- | --- | --- | --- | --- |
| DICER1 | Rabbit | Bethyl Laboratories | 1:5000 | A301-936A |
| AGO2 (eIF2C2) | Mouse Monoclonal | Abnova | 1:1000 | H00027161-M01 |
| HA | Rat Monoclonal ( clone 3F10) | Roche | 1:1000 | 11 867 423 001 |
| RCK/p54 | Rabbit | Bethyl Laboratories | 1:10000 | A300-461A |
| HRS | Rabbit | Bethyl Laboratories | 1:1000 | A300-989A |
| Calnexin | Rabbit | Cell Signaling Technology | 1:1000 | #2679S |
| Alix | Mouse Monoclonal | Santa Cruz | 1:500 | sc53538 |
| HuR | Mouse Monoclonal (clone 3A2) | Santa Cruz | 1:1000 | sc-5261 |
| eIF2 alpha | Rabbit | Cell Signaling Technology | 1:500 | #9722 |
| Phospho eIF2 alpha | Rabbit | Cell Signaling Technology | 1:500 | #9721 |
| Cleaved PARP | Rabbit | Cell Signaling Technology | 1:1000 | #9541S |
| Rab5 | Rabbit | Cell Signaling Technology | 1:1000 | #3547P |
| Rab7 | Rabbit | Cell Signaling Technology | 1:1000 | #9367 |
| XRN1 | Rabbit | Bethyl Laboratories | 1:10000 | A300-443A |
| ADRP | Mouse | Santa Cruz | 1:200 | sc-377429 |
| HSP90 | Rabbit | Cell Signaling Technology | 1:1000 | #4877 |
| CD63 | Mouse | BD Pharmingen | 1:500 | 556019 |
| GAPDH | Mouse | Sigma | 1:1000 | G8795 |
| β-tubulin | Mouse | Sigma | 1:1000 | T5201 |
| β-actin (HRP conjugated) | Mouse | Sigma | 1:10000 | A3854-200UL |

**Supplementary Table 2.** List of secondary antibodies used

| Secondary Antibody | Source | Dilution for Western blot | Catalog no. |
| --- | --- | --- | --- |
| HRP Goat anti-mouse | Life Technologies | 1:8000 | 626520 |
| HRP Goat anti-Rabbit | Life Technologies | 1:8000 | 656120 |
| HRP Goat anti-Rat | Life Technologies | 1:8000 | 629520 |

**Supplementary Table 3.** List of fluorochrome tagged secondary antibodies used:

| Secondary Antibody | Source | Dilution for Microscopy | Catalog no. |
| --- | --- | --- | --- |
| Anti-rabbit Alexa Fluor® 568 | Molecular Probes | 1:500 | A11036 |
| Anti-rabbit Alexa Fluor® 647 | Molecular Probes | 1:500 | A21245 |
| Anti-rat Alexa Fluor® 568 | Molecular Probes | 1:500 | A11077 |

**Supplementary Table 4.**
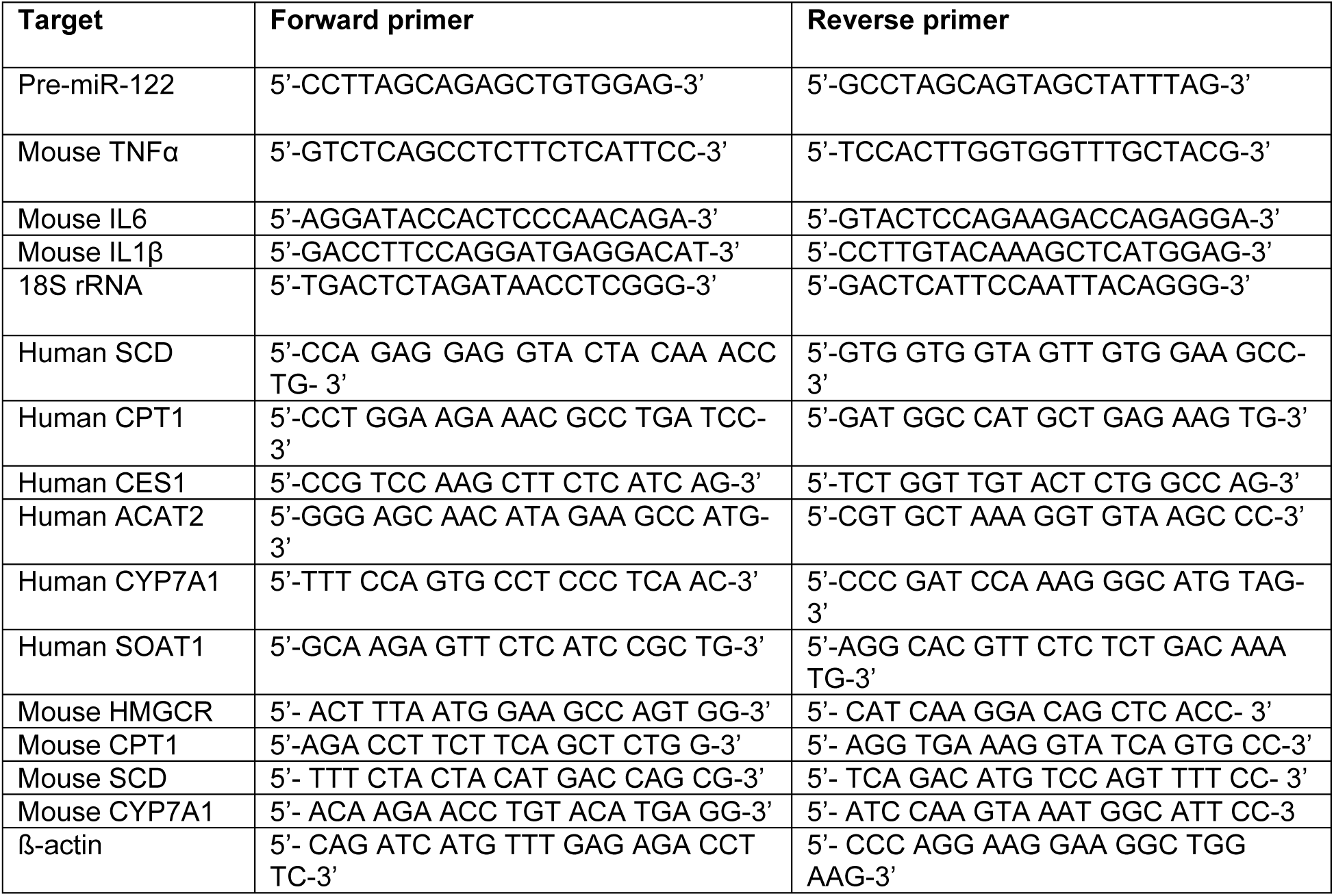
List of primers used for RT-qPCR detection of mRNAs

**Supplementary Table 5.** List of primers used for RT-qPCR detection of miRNAs:

| Target miRNA | Assay ID |
| --- | --- |
| Human miR-122 | 000445 |
| Human miR-122* | 002130 |
| Human miR-16 | 000391 |
| Mouse miR-155 | 002571 |
| Human miR-24 | 000402 |
| Human miR-21 | 000397 |
| Human miR-33a | 002135 |
| Human miR-29a | 000412 |
| Human let-7a | 000377 |
| U6 snRNA | 001973 |

**Supplementary Table 6.**
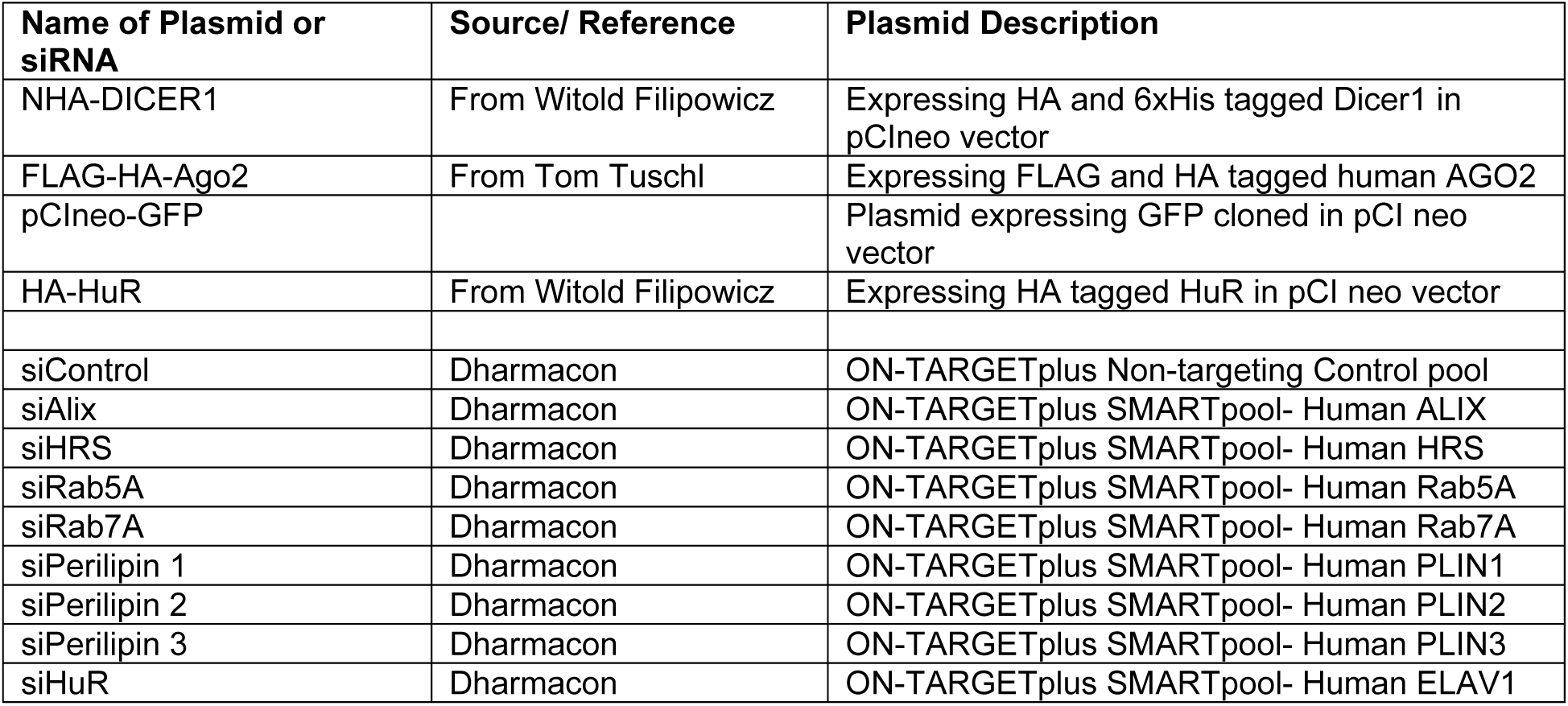
List of Plasmids and siRNAs used

## Appendix Figures

**Appendix Figure 1:**
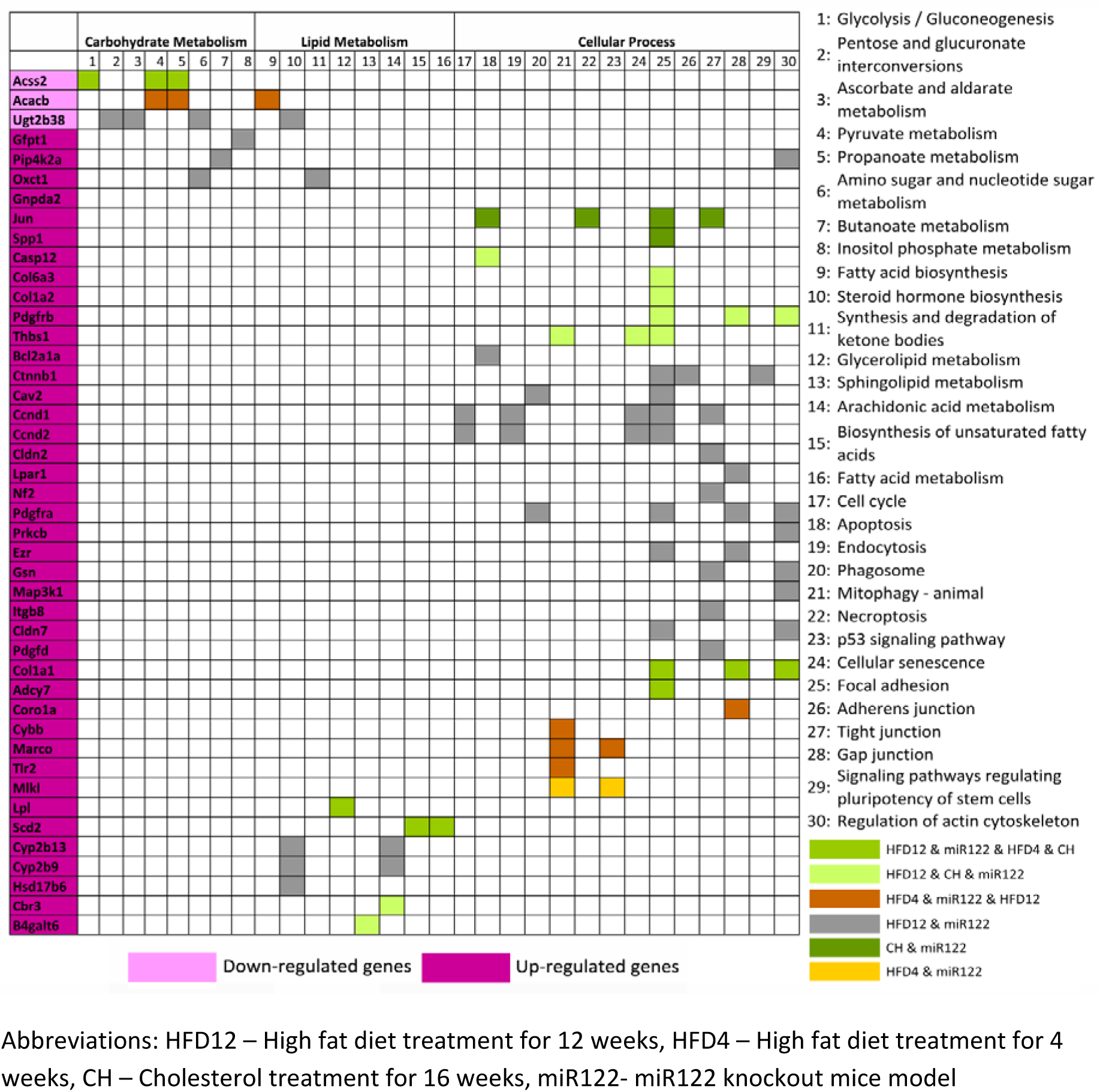
High fat exposure related hepatic cell stress results in deregulation of miR- 122 target genes involved in cellular metabolism and cellular process associated pathways. Differentially expressed miR-122 target genes that occur upon high fat exposure in hepatic cells participate in multiple pathways belonging to carbohydrate metabolism, lipid metabolism or cellular process as elucidated here. Abbreviations: HFD12 – High fat diet treatment for 12 weeks, HFD4 – High fat diet treatment for 4 weeks, CH – Cholesterol treatment for 16 weeks, miR122- miR122 knockout mice model.

**Appendix Figure 2:**
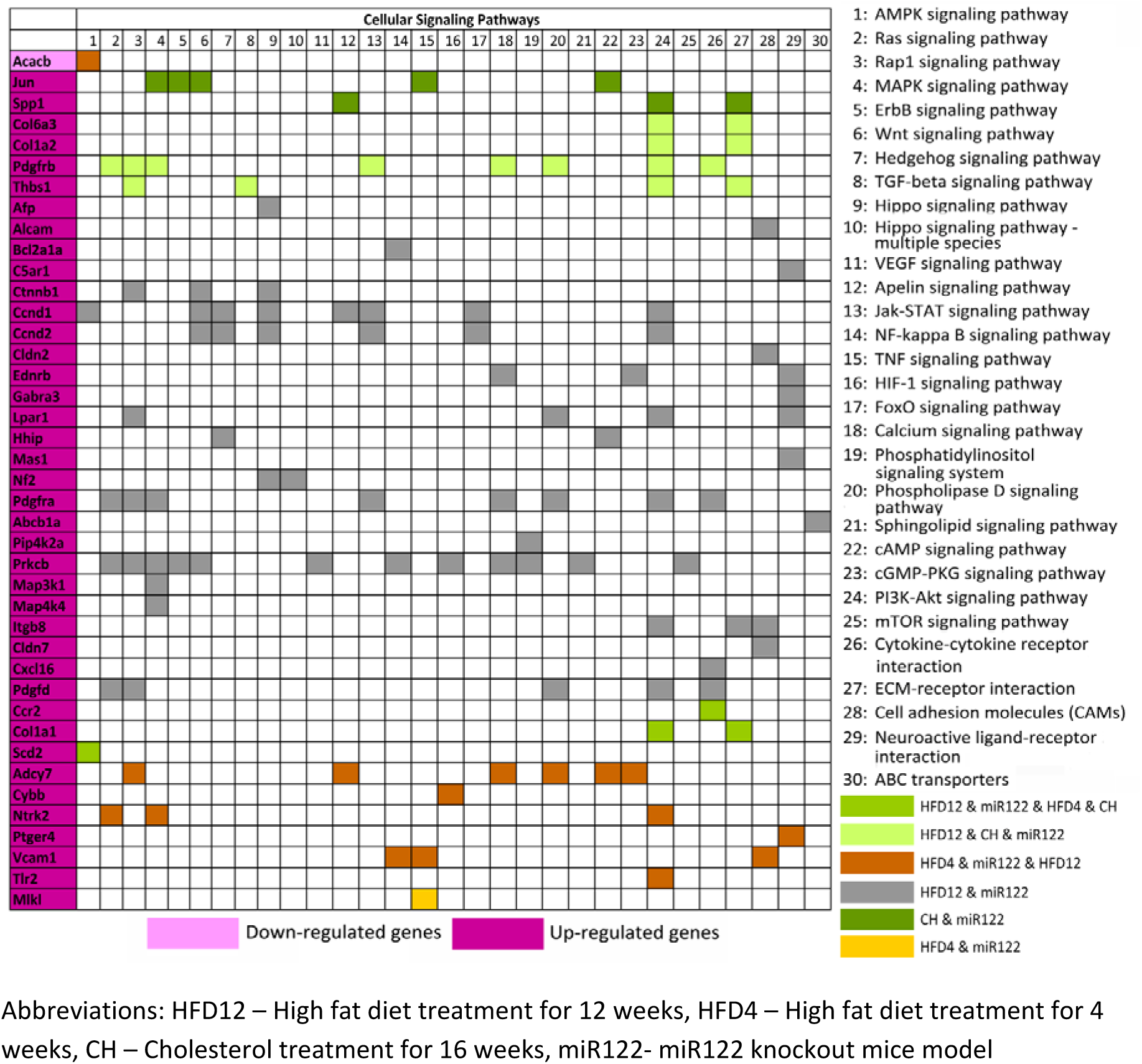
Chronic high fat exposure associated stress in hepatic cells leads to the deregulation of miR-122 target genes involved in multiple cellular signalling pathways. Pathway mapping of differentially expressed miR-122 target genes that occur upon prolonged high fat diet treatment in mice models elucidated that multiple proteins encoded by these genes participate in a range of signalling pathways as exemplified here. Abbreviations: HFD12 – High fat diet treatment for 12 weeks, HFD4 – High fat diet treatment for 4 weeks, CH – Cholesterol treatment for 16 weeks, miR122- miR122 knockout mice model.

